# Development and validation of a deep learning framework for Alzheimer’s disease classification

**DOI:** 10.1101/832519

**Authors:** Vijaya B. Kolachalama, Shangran Qiu, Prajakta S. Joshi, Matthew I. Miller, Chonghua Xue, Cody Karjadi, Gary H. Chang, Anant S. Joshi, Brigid Dwyer, Shuhan Zhu, Michelle Kaku, Yan Zhou, Yazan J. Alderazi, Arun Swaminathan, Sachin Kedar, Marie-Helene Saint-Hilaire, Sanford H. Auerbach, Jing Yuan, E. Alton Sartor, Rhoda Au

**Author notes:** **Corresponding author**: Vijaya B. Kolachalama, PhD, 72 E. Concord Street, Evans 636, Boston, MA - 02118.

## Abstract

Alzheimer’s disease (AD) is the primary cause of dementia worldwide (*1*), with an increasing morbidity burden that may outstrip diagnosis and management capacity as the population ages. Current methods integrate patient history, neuropsychological testing and magnetic resonance imaging (MRI) to identify likely cases, yet effective practices remain variably-applied and lacking in sensitivity and specificity (*2*). Here we report an explainable deep learning strategy that delineates unique AD signatures from multimodal inputs of MRI, age, gender, and mini-mental state examination (MMSE) score. Our framework linked a fully convolutional network (FCN) to a multilayer perceptron (MLP) to construct high resolution maps of disease probability from local brain structure. This enabled precise, intuitive visualization of individual AD risk en route to accurate diagnosis. The model was trained using clinically-diagnosed AD and cognitively normal (NC) subjects from the Alzheimer’s Disease Neuroimaging Initiative (ADNI) dataset (n=417) (*3*), and validated on three independent cohorts: the Australian Imaging, Biomarker & Lifestyle Flagship Study of Ageing (AIBL, n=382) (*4*), the Framingham Heart Study (FHS, n=102) (*5*), and the National Alzheimer’s Coordinating Center (NACC, n=582) (*6*). Model performance was consistent across datasets, with mean accuracy values of 0.966, 0.948, 0.815, and 0.916 for ADNI, AIBL, FHS and NACC, respectively. Moreover, our approach exceeded the diagnostic performance of a multi-institutional team of practicing neurologists (n=11), and high-risk cerebral regions predicted by the model closely tracked postmortem histopathological findings. This framework provides a clinically-adaptable strategy for using routinely available imaging techniques such as MRI to generate nuanced neuroimaging signatures for AD diagnosis, as well as a generalizable approach for linking deep learning to pathophysiological processes in human disease.

## Introduction

Millions worldwide continue to suffer from AD, while attempts to develop effective disease modifying treatments remain stalled. Though tremendous progress has been made towards detecting AD pathology using cerebrospinal fluid (CSF) biomarkers (*7–9*), as well as positron emission tomography (PET) amyloid (*10, 11*), and tau (τ) imaging (*12, 13*), these modalities often remain limited to research contexts. Instead, current standards of diagnosis depend on highly skilled neurologists to conduct an examination that includes inquiry of patient history, an objective cognitive assessment such as bedside MMSE or neuropsychological testing (*2*), and a structural MRI to rule in findings suggestive of AD (*7*). Clinicopathological studies suggest the diagnostic sensitivity of clinicians range between 70.9-87.3% and specificity between 44.3%- 70.8% (*14*). While MRIs reveal characteristic cerebral changes noted in AD such as hippocampal and parietal lobe atrophy (*15*), these characteristics are considered to lack specificity for imaging-based AD diagnosis (*7, 16–18*). Given this relatively imprecise diagnostic landscape, as well as the invasive nature of CSF and PET diagnostics and a paucity of clinicians with sufficient AD diagnostic expertise, advanced machine learning paradigms such as deep learning (*19–21*), offer ways to derive high accuracy predictions from MRI data collected within the bounds of neurology practice.

Recent studies demonstrate the application of deep learning approaches such as convolutional neural networks (CNNs) for MRI and multimodal data-based classification of cognitive status (*22*). Despite the promising results, these models have yet to achieve full integration into clinical practice for several reasons. First, there is a lack of external validation of deep learning algorithms since most models are trained and tested on a single cohort. Second, there is a growing notion in the biomedical community that deep learning models are “black-box” algorithms (*23*). In other words, although deep learning models demonstrate high-accuracy classification across a broad spectrum of disease, they neither elucidate the underlying diagnostic decisions nor indicate the input features associated with the output predictions. Lastly, given the uncertain onset and heterogeneity of symptoms seen in AD, a computerized individual-level characterization of AD remains unresolved. Considering these factors, we surmise that the clinical potential of deep learning is attenuated by a lack of external validation of single cohort-driven models, and an increasing use of opaque decision-making frameworks. Thus, overcoming these challenges is not only crucial to harness the potential of deep learning algorithms to improve patient care, but to also pave the way for explainable evidence-based machine learning in the medical imaging community. To address these limitations, we developed a novel deep learning framework that links a fully convolutional network (FCN) to a traditional multilayer perceptron (MLP) to generate high-resolution visualizations of AD risk that can then be used for accurate predictions of AD status (Fig. 1). Four distinct datasets were chosen for model development and validation (Fig. 2, Table 1). Association of model predictions with neuropathological findings along with a head-to-head comparison of the model performance with a team of neurologists underscored the validity of the deep learning framework.

**Figure 1:**
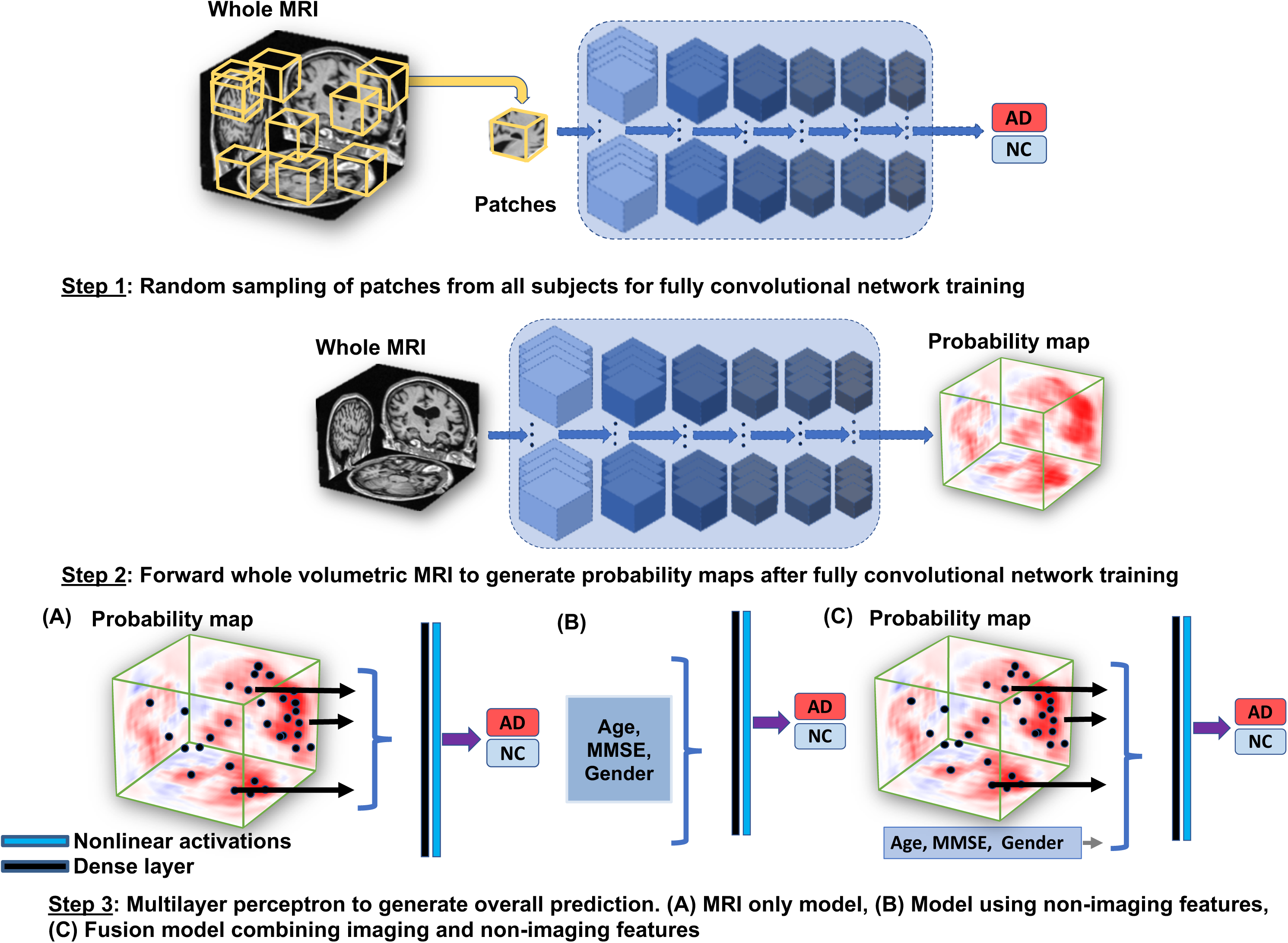
Schematic of the deep learning framework. The fully convolutional network (FCN) model was developed using a patch-based strategy in which randomly selected samples (sub-volumes of size 47×47×47 voxels) of T1-weighted full MRI volumes were passed to the model for training (Step 1). The corresponding AD status of the individual served as the output for the classification model. Given that the operation of FCNs is independent of input data size, the model led to the generation of participant-specific disease probability maps (DPMs) of the brain (Step 2). Selected voxels of high-risk from the DPMs were then passed to the MLP for binary classification of disease status (Model A in Step 3). As a further control, we used only the non-imaging features including age, gender and MMSE and developed an MLP model to classify individuals with AD and the ones with normal cognition (Model B in Step 3). We also developed another model that integrated multimodal input data including the selected voxels of high-risk DPMs alongside age, gender and MMSE score to perform binary classification of AD status (Model C in Step 3).

**Figure 2:**
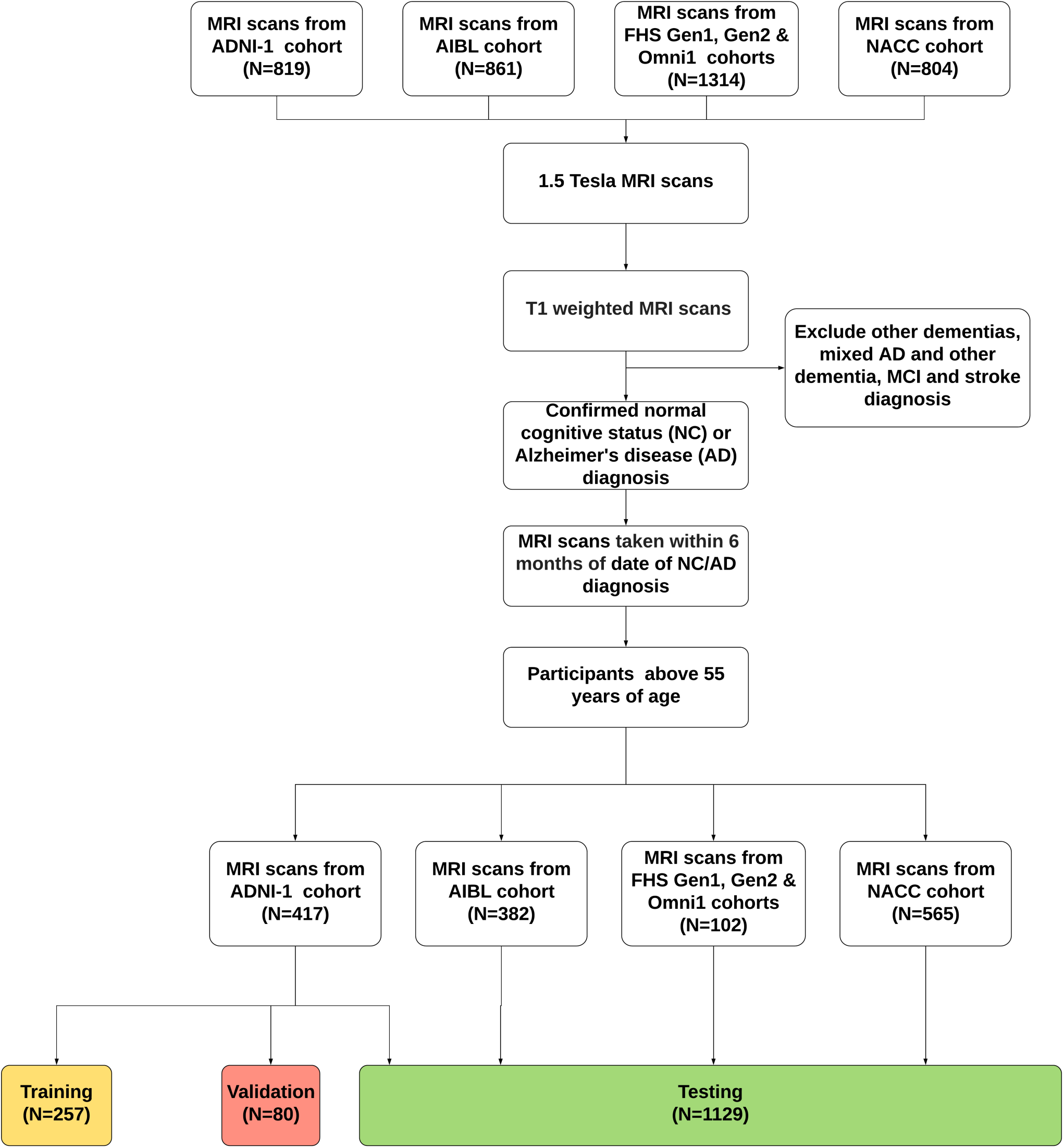
Subject selection criterion. In each dataset, T1-weighted, 1.5 Tesla MRIs were selected from participants (See Methods for more details). Only MRIs gathered within 6 months of AD diagnosis or last confirmed clinical visit (in the case of NC participants) were included for analysis. From there, ADNI data was split in a 2:1:1 ratio for training, validation, and testing sets, and fully trained models were applied to NACC, FHS, and AIBL to assess model generalizability.

**Table 1:** Study population and characteristics. Four independent datasets including (a) the Alzheimer’s Disease Neuroimaging Initiative (ADNI) dataset, (b) the Australian Imaging, Biomarker & Lifestyle Flagship Study of Ageing (AIBL), (c) the Framingham Heart Study (FHS), and (d) the National Alzheimer’s Coordinating Center (NACC)) were used for this study. The ADNI dataset was randomly split in the ratio of 2:1:1, where 50% of it was used for model training, 25% of the data was used for internal validation and the rest was used for internal testing. The best performing model on the validation dataset was selected for making predictions on the ADNI test data as well as on the AIBL, FHS and NACC datasets, which served as external test datasets for model validation. All the MRI scans considered for this study were performed on individuals within ±6 months from the date of clinical diagnosis.

| Dataset | ADNI (a) |  |  | AIBL (b) |  |  | FHS (c) |  |  | NACC (d) |  |  |
| --- | --- | --- | --- | --- | --- | --- | --- | --- | --- | --- | --- | --- |
| Characteristic | NC<br>(n=229) | AD<br>(n=188) | p-value | NC<br>(n=320) | AD<br>(n=62) | p-value | NC<br>(n=73) | AD<br>(n=29) | p-value | NC<br>(n=356) | AD<br>(n=209) | p-value |
| Age, y, median<br>[range] | 76<br>[60, 90] | 76<br>[55, 91] | 0.4185 | 72<br>[60, 92] | 73<br>[55, 93] | 0.5395 | 73<br>[57, 100] | 81<br>[67, 94] | <0.0001 | 74<br>[56, 94] | 77<br>[55, 95] | 0.0332 |
| Education, y, median<br>[range] | 16<br>[6, 20] | 16<br>[4, 20] | <0.0001 | N.A. | N.A. | N.A. | 14<br>[8, 25] | 13<br>[5, 25] | 0.3835 | 16<br>[0, 22]* | 14.5<br>[2, 24]# | 0.8363 |
| Gender, male (%) | 119<br>(51.96) | 101<br>(53.72) | 0.7677 | 144<br>(45.00) | 24<br>(38.71) | 0.4031 | 37<br>(50.68) | 12<br>(41.38) | 0.5105 | 126<br>(35.39) | 95<br>(45.45) | 0.0203 |
| MMSE, median<br>[range] | 29<br>[25, 30] | 23.5<br>[18, 28] | <0.0001 | 29<br>[25, 30] | 21<br>[6, 28] | <0.0001 | 29<br>[22, 30]⊥ | 25<br>[10, 29] | <0.0001 | 29<br>[20, 30]↑ | 22<br>[0, 30]‡ | <0.0001 |
| APOE4, positive (%) | 61 (27) | 124 (66) | <0.0001 | 11 (3.4) | 12 (19.4) | <0.0001 | 13<br>(17.81) | 11<br>(40.74)∧ | 0.0355 | 102<br>(28.65) | 112<br>(53.59) | <0.0001 |

\* -- Education information not available for two subjects # -- Education information not available for one subject ↑ -- MMSE score was not available for one subject ‡ -- MMSE score was not available for one subject ⊥ -- Six subjects do not have available MMSE scores ∧ -- APOE4 information not available for one subject
\* -- Education information not available for two subjects

## Results

Our dual deep learning pipeline can link an FCN to an MLP to predict AD status directly from MRI data or from a combination of MRI data and readily available non-imaging data (Fig. 1). The FCN portion of the framework generated high-resolution visualizations of overall AD risk in individuals as a function of local cerebral morphology. We refer to these visualizations as disease probability maps (DPMs). The MLP then used DPMs directly (Model A in Fig. 1), or a set of non-imaging features such as age, gender and MMSE score (Model B in Fig. 1), or a multimodal input data comprising DPMs, MMSE score, age and gender (Model C in Fig. 1), to accurately predict AD status across four independent cohorts (Table 1, Fig. 2). We chose these known AD risk factors because they can be easily obtained by non-AD specialists. The FCN was trained to predict disease probability from randomly-selected patches (sub-volumes) of pixels sampled from the full MRI volume (Fig. 1 & Table S1). Given that this type of network accepts input of arbitrary size, application of the sub-volumetrically trained FCN could then be used to construct high resolution DPMs without the need to redundantly decompose full-sized test images.

Rapid processing of individual MRI volumes generated volumetric distributions of local AD probabilities in the brains of affected and unaffected individuals, respectively (Figs. 3a, S1-S3). In order to assess the anatomical consistency of AD suggestive morphology hot spots derived from these distributions, population-wide maps of Matthew’s Correlation Coefficient (MCC) were constructed. MCC mapping enabled identification of areas from which correct predictions of disease status were most frequently derived (Fig. 3b), thus acting as a means to demonstrate structures most affected by neuropathological changes in AD.

**Figure 3:**
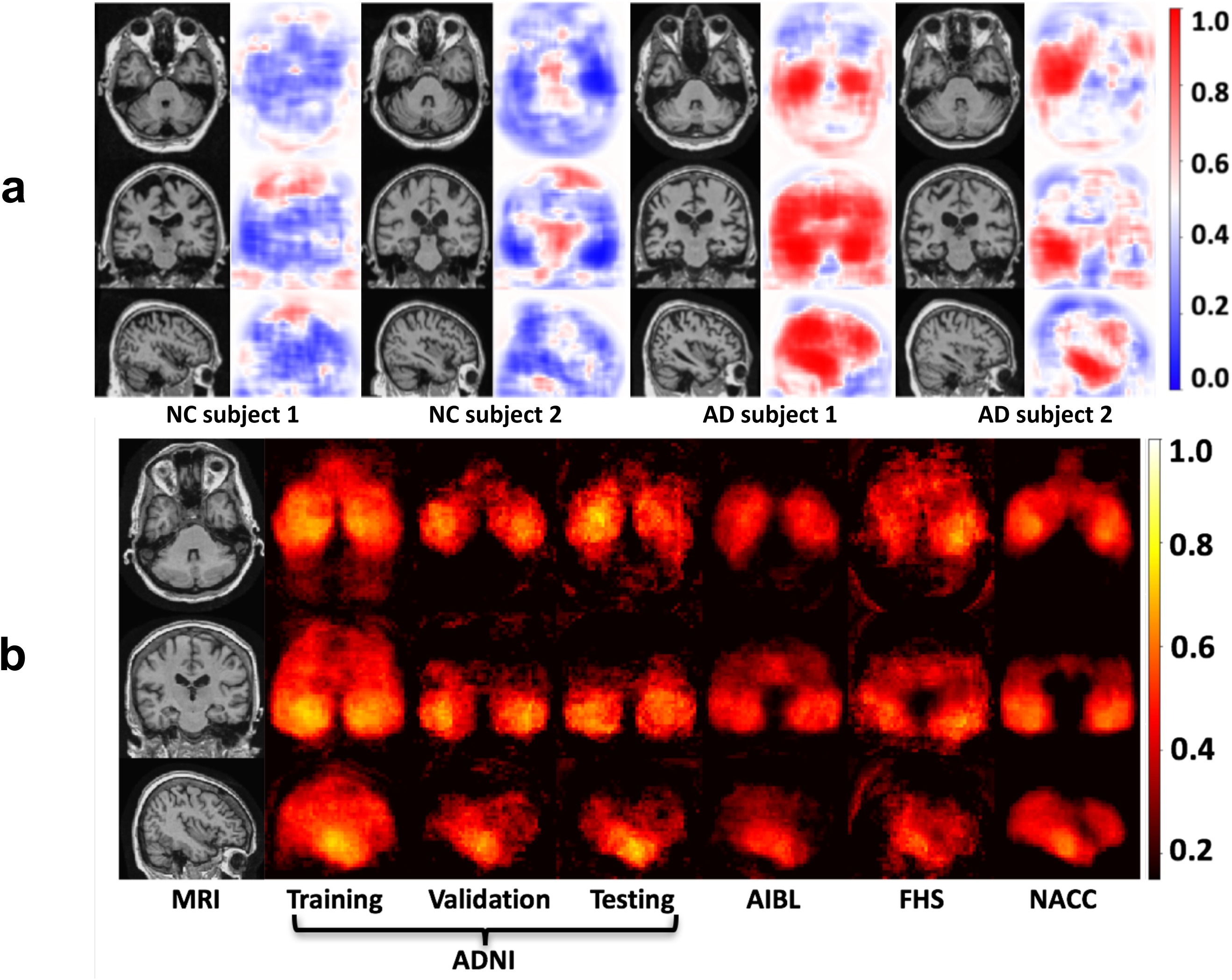
Fully convolutional network (FCN) model performance. (a) Disease probability maps (DPMs) generated by the FCN model highlight high-risk brain regions that are associated with AD pathology. Individual cases are shown (two NC and two AD), where the blue color indicates low-risk and red indicates high-risk of AD. The first two individuals were clinically confirmed to have normal cognition whereas the other two individuals had clinical diagnosis of AD. (b) Pixel-wise maps of Matthew’s correlation coefficient were computed independently across all the datasets to demonstrate predictive performance derived from all regions within the brain.

As confirmation, average regional probabilities extracted from selected segmented brain regions (Fig. 4), were highly associated with AD positive findings reported in postmortem neuropathology examinations. Specifically, these regions correlated with the locations and numerical frequency of Aβ and τ pathologies reported in available autopsy reports from the FHS dataset (n=11) (Table S2). Postmortem data indicated that, in addition to predicting higher region-specific AD probabilities in individuals with disease compared to those without, proteinopathies were more frequent in cerebral regions implicated by the model in AD (Fig. 4). Model-predicted regions of high AD risk overlapped with the segmented regions that were indicated to have high localized deposition of Aβ and τ. Additionally, predicted AD risk within these zones increased with pathology scores. Given that these postmortem findings are definitive in terms of confirming AD, these physical findings grounded our computational predictions in biological evidence.

**Figure 4:**
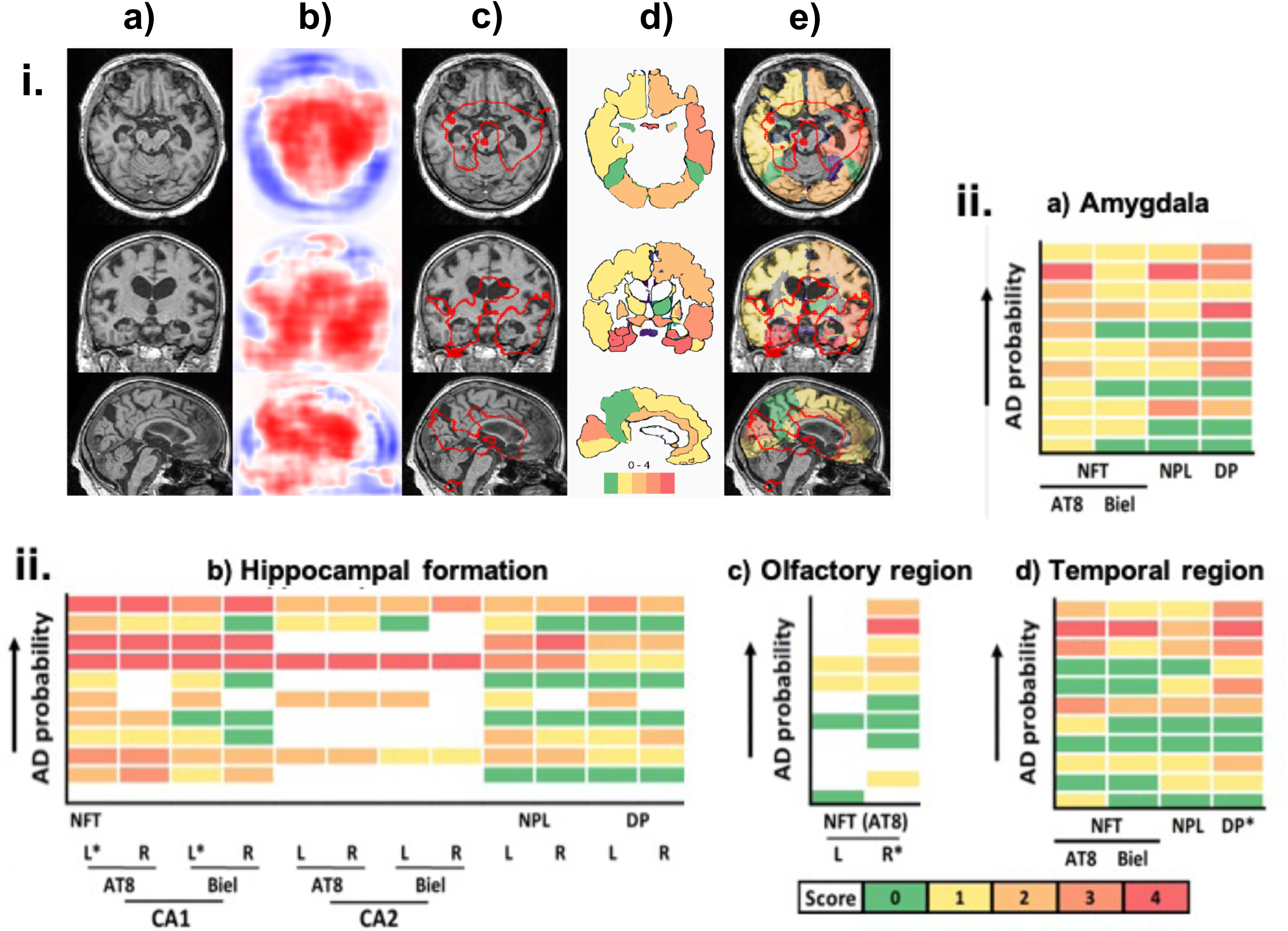
Correlation of model findings with neuropathology. Overlap of model predicted regions of high AD risk with postmortem findings of AD pathology. The scans were generated by averaging across 11 individuals from the FHS cohort on whom the postmortem findings were also available. (i) The first column (a) shows MRI slices in three different planes followed by a column (b), which shows corresponding model predicted DPMs. A cutoff value of 0.85 was chosen to delineate the regions of high AD risk and overlapped with the MRI scan in the next column (c). The next column (d), depicts a segmented mask of cortical and sub-cortical structures of the brain obtained from FreeSurfer (*28*). A sequential color-coding scheme denotes different levels of pathology ranging from green (0-low) to pale red (4-high). The final column (e), shows the overlay of the MR scan, DPMs of high AD risk and the color-coded regions based on pathology grade. (ii) We then qualitatively assessed trends of neuropathological findings from the FHS dataset (n=11). The same color-coding scheme as described above was used to represent the pathology grade (0-4) in the heat maps. The boxes colored in ‘white’ in the heat maps indicate missing data. Using the Spearman’s Rank correlation coefficient test, an increasing AD probability risk was associated with a higher grade of Aβ and τ accumulation, in the hippocampal formation, the temporal region, and the olfactory bulbs, respectively. A similar trend was qualitatively observed in the amygdala; however, this region did not reach statistical significance at the 0.05 level.

Furthermore, DPMs provided an information-dense feature that yielded sensitive and specific binary predictions of AD status when passed independently to the MLP portion of the framework (Model A in Figs. 5a-5b). An MLP trained using just the non-imaging features such as age, gender and MMSE score also was predictive of AD status (Model B in Figs. 5a-5b). Model performance was further improved by expanding the MLP input to include DPMs, gender, age, and MMSE score (Model C in Figs. 5a-5b). When other non-imaging features such as ApoE status were included, model performance slightly improved (Fig. S4 & Table S3). Given the proportionality between age and global cerebral atrophy (*17, 18*), addition of non-imaging variables at the MLP stage also allowed us to control for the natural progression of cerebral morphological changes over the lifespan.

**Figure 5:**
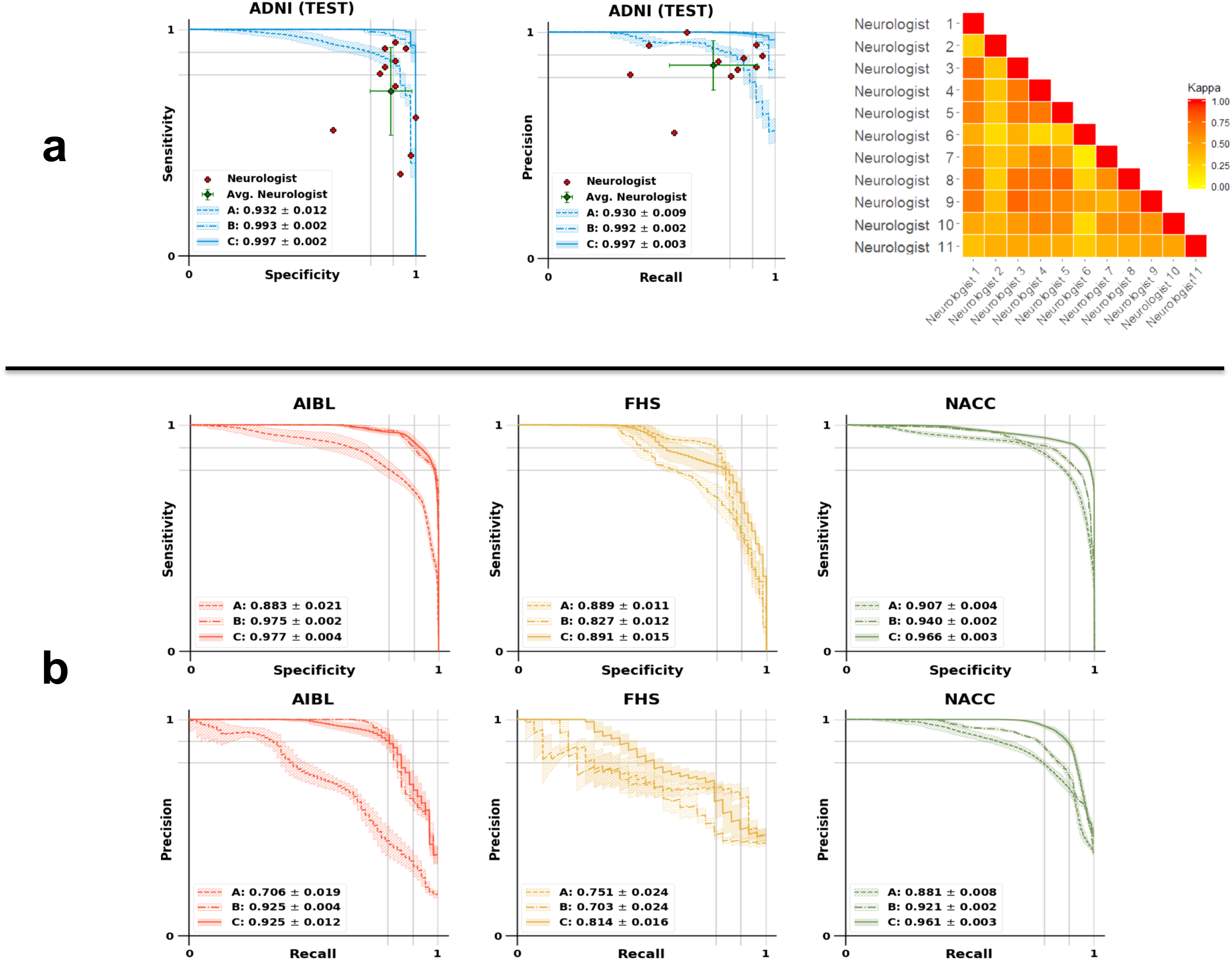
Performance of the MLP model for AD classification and model comparison with neurologists. (a) Sensitivity-specificity (SS) and precision-recall (PR) curves showing the sensitivity, the true positive rate, versus specificity, the true negative rate, calculated on the ADNI test set. Individual neurologist performance is indicated by the red (+) symbol and averaged neurologist performance along with the error bars is indicated by the green (+) symbol on both the SS and PR curves on the ADNI test data. Visual description of pairwise Cohen’s kappa (K), which denotes the inter-operator agreement between all the 11 neurologists is also shown. (b) SS and PR curves calculated on the AIBL, FHS and NACC datasets, respectively. For all cases, Model A indicates the performance of the MLP model that used MRI data as the sole input, model B is the MLP model with non-imaging features as input and model C indicates the MLP model that used MRI data along with age, gender and MMSE values as the inputs for binary classification.

We also compared performance of the deep learning models against an international group of clinical neurologists recruited to provide impressions of disease status from a randomly-sampled cohort of ADNI participants whose MRI, MMSE score, age and gender were provided. The performance of the neurologists (Figs. 5a-5b), indicated variability across different clinical practices, with a moderate inter-rater agreement as assessed by pairwise kappa (K) scoring (Fig. 5a; average K=0.493±0.16). Interestingly, we noted that the deep learning model that was based on MRI data alone (Model A; Accuracy: 0.858±0.016; Table S4), outperformed the average neurologist (Accuracy: 0.823±0.094; Table S5). When age, gender and MMSE information was added to the model, then the performance increased significantly (Model C; Accuracy: 0.963±0.013; Table S4). High classification performance of the deep learning model was confirmed using other metrics (Table S4), and sub-group analyses (Fig. S5). Also, we examined the model performance visually by respective clustering of AD cases and the ones who had normal cognition (NC) in a t-distributed stochastic neighbor embedding (t-SNE) (*26*), which employed features from the final hidden layer of the MLP (Fig. 6). The t-SNE method takes high-dimensional data and creates a low-dimensional representation of that data, so that it can be easily visualized.

**Figure 6:**
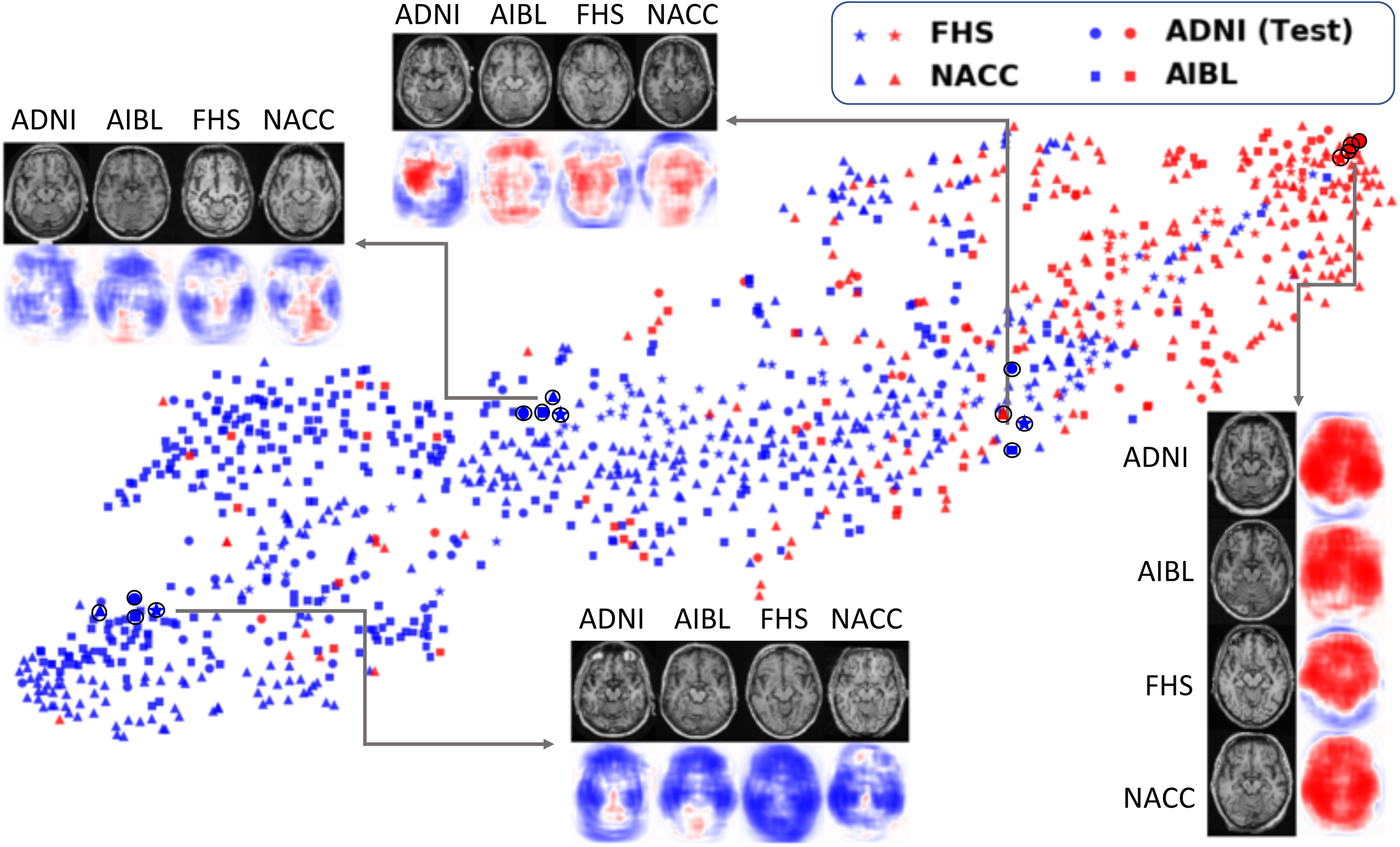
Visualization of data. FCN-based outputs that served as input features to the MLP model were embedded in a two-dimensional plot generated using t-SNE, a method for visualizing high-dimensional data, for the two classes (AD and NC). The color (blue versus red) was used to distinguish NC from AD cases, whereas a unique symbol shape was used to represent individuals derived from the same cohort. Several individual cases that were clinically confirmed to have AD or normal cognition are also shown (indicated as a black circle overlaying the respective data point in the figure). The plot also indicates co-localization of subjects in the feature space based on the disease state and not on the dataset of origin.

It is worth noting that our strategy represents a significant increase in computational efficiency over a traditional CNN approach to the same task (Step 1 in Fig. 1 vs Fig. S6). Given fixed dense layer dimensions, generation of DPMs from traditional CNNs requires not only sub-volumetric training, but also sub-volumetric application to full-sized MRI volumes (Table S1 vs Table S6), obligating repeated computations in order to calculate local probabilities of disease status. By circumventing this rigidity, our approach readily generates DPMs (Step 2 in Fig. 1), which can be integrated with multimodal clinical data for AD diagnosis (Step 3 in Fig. 1). As such, this work extends recently reported efforts to abstract visual representations of disease risk directly from medical images (*24*), and also represents the first application of FCNs to disease classification tasks as opposed to semantic segmentation (*25*). In a head-to-head comparison, the FCN model outperformed the traditional CNN model in predicting AD status, and this result was consistent across all the datasets (Fig. S7 & Table S7).

## Discussion

Our deep learning framework links an FCN to an MLP and generates high resolution DPMs for neurologist-level diagnostic accuracy of AD status. The intuitive local probabilities outputted by our model are readily interpretable, thus contributing to the growing movement towards explainable artificial intelligence in medicine, and deriving an individualized phenotype of insidious disease from conventional diagnostic tools. Indeed, the DPMs provide a means for tracking conspicuous brain regions implicated in AD during diagnosis. We then aggregated DPMs across the entire cohort to demonstrate population-level differences in neuroanatomical risk mapping of AD and NC cases. Critically, by the standards of several different metrics, our model displayed high predictive performance, yielding high and consistent values on all the test datasets. Such consistency between cohorts featuring broad variance in MRI protocol, geographic location, and recruitment criteria, suggests a strong degree of generalizability. Thus, these findings demonstrate innovation at the nexus of medicine and computing, simultaneously contributing new insights to the field of computer vision while also expanding the scope of biomedical applications of neural networks.

DPMs were created by element-wise application of a softmax function to the final array of activations generated by the FCN. This step enabled the conversion of abstract tensor encodings of neuroanatomical information to probability arrays demonstrating the likelihood of AD at different locations in the brain given their local geometry. Alternatively put, the model develops a granular conceptualization of AD-suggestive morphologies throughout the brain, and then uses this learning information in test cases to assess the probability of AD-related pathophysiologic processes occurring at each region. The simple presentation of these probabilities as a coherent colormap displayed alongside traditional neuroimaging thus allows a point-by-point prediction of where disease-related changes are likely to be present. While DPMs are generated using the FCN framework, a conceptual understanding of neural networks is not warranted for their interpretation. Recent work has also demonstrated effective differentiation of AD and NC cases using a patch-based sampling algorithm (*27*), but is limited by simultaneous reliance on MRI and FDG-PET as well as a model whose inputs are computed as scalar averages of intensities from multi-voxel cerebral loci. Furthermore, we believe that the broader notion of disease process mapping with deep learning has the potential to be applied in many fields of medicine. The simple presentation of disease risk as a coherent colormap overlaid on traditional imaging modalities aids interpretability. This is in contrast to saliency mapping strategies that highlight certain pixels based only on their utility to the internal functioning of a network (*25*), as well as methods that highlight penultimate-layer activation values (*27*). Consequently, informative anatomical information is abstracted and lost. Our work builds upon such advances by requiring just a single imaging modality en route to mapping an array of raw pixel values to a DPM that isomorphically preserves neuroanatomical information. While traditional neural networks require an input of fixed magnitude, FCNs are capable of acting on inputs of arbitrary size. Thus, the same patch-trained model was then applied to test inputs of full MRIs. This allowed the local representations of AD status learned by the model during training to be transferred to the task of predicting disease processes throughout the entire brain. Importantly, just as the input size was larger in testing, so too was the size of the output. Thus, during application of the FCN to unseen MRI stacks, the model produces two 3D matrices of activation values, which get converted to arrays of probability values by element-wise softmax application. Together, these two arrays represented spatial distributions of AD and NC probabilities, respectively.

Certainly, limitations to the current study must be acknowledged. We considered a case-control population, where two sub-populations were chosen in advance that were either cognitively normal (NC) or have the diagnosis (AD). Therefore, the clinical relevance of our model is restricted to scenarios when patients are screened out for other forms of dementia or pseudodementia. While this may not be the clinical scenario faced by the majority of neurologists, who see individuals with any number of diagnoses and then must distinguish between them, certainly it is still an interesting and important clinical question. Furthermore, the circularity that MMSE tests and other demographic parameters used to determine clinical diagnosis may capture a biased diagnosis which in turn may differ across the different studies. Consequently, our results indicate the potential to augment standard methods of AD management via emerging computational strategies. It is also worthwhile to note that non-imaging data-based models performed better on AIBL and NACC data, while the MRI-based model performed better on the FHS data. As such, the MMSE value was a key element in the study criteria for ADNI, AIBL and NACC, and this may explain why the non-imaging data-based model performed better on these datasets. Since FHS is a community cohort, it more or less remained as a fairly unbiased dataset for analysis. Despite this study selection, our MRI-based model provides compelling evidence that use of an imaging biomarker alone can provide strong classification in a deep learning framework.

Our approach has significant translational potential beyond AD diagnosis. Indeed, the tissue-level changes predicted by our model suggest the prospect of directly highlighting areas of pathophysiology across a spectrum of human disease. This ability may be particularly useful in conditions where diffuse symptomologies are accompanied by insidious lesions, including non-AD dementias and cognitive conditions. Additionally, it may be of interest in future studies to determine whether the well-defined pattern of high-risk findings from the currently-presented framework may follow regions of interest from PET. In such cases, our model may aid in both noninvasive monitoring of AD development and expanding access to high quality care in resource-limited settings.

In conclusion, our deep learning framework was able to obtain high accuracy AD classification signatures from MRI data, and our model was validated against data from independent cohorts, neuropathological findings and expert-driven assessment. If confirmed in clinical settings, this approach has the potential to expand the scope of neuroimaging techniques for disease detection and management. Further validation could lead to improved care and outcomes compared with current neurologic assessment, as the search for disease modifying therapies continues.

## Methods

### Study participants and data collection

Data from ADNI, AIBL, FHS, and NACC cohorts were used in the study (Fig. 2 & Table 1). ADNI is a longitudinal multicenter study designed to develop clinical, imaging, genetic, and biochemical biomarkers for the early detection and tracking of AD (*3*). AIBL, launched in 2006, is the largest study of its kind in Australia and aims to discover biomarkers, cognitive characteristics, and lifestyle factors that influence the development of symptomatic AD (*4*). FHS is a longitudinal community cohort study and has collected a broad spectrum of clinical data from three generations (*5*). Since 1976, FHS expanded to evaluate factors contributing to cognitive decline, dementia, and AD. Finally, NACC that was established in 1999, maintains a large relational database of standardized clinical and neuropathological research data collected from AD centers across the US (*6*).

Model training, internal validation and testing were performed on the ADNI dataset. Following training and internal testing on the ADNI data, we validated the predictions on AIBL, FHS, and NACC. The criterion for selection included individuals over age ≥55 years, with 1.5 Tesla, T1-weighted MRI scans taken within ±6 months from the date of clinically confirmed diagnosis of AD or NC (Fig. 2). We excluded cases including AD with mixed dementia, non-AD dementias, history of severe traumatic brain injury, severe depression, stroke, and brain tumors, as well as incident major systemic illnesses. Note that this inclusion and exclusion criterion was adapted from the baseline recruitment protocol developed by the ADNI study (*3*), and to maintain consistency, the same criterion was applied to other cohorts as applicable. This led to the selection of 417 individuals from the ADNI cohort 382 individuals from AIBL, 102 FHS participants, and 565 individuals from the NACC cohort. If an individual had multiple MRI scans taken within the time window, then we selected the scan closest to the date of clinical diagnosis. For all these selected cases, age, gender and MMSE score were available.

### Algorithm development

A fully convolutional network (FCN) was designed to input a registered volumetric MRI scan of size 181×217×181 voxels and output the AD class probability at every location. We employed a novel, computationally efficient patch-wise training strategy to train the FCN model (Fig. 1). This process involved random sampling of 3000 volumetric patches of size 47×47×47 voxels from each training subject’s MRI scan and used this information to predict the output of interest. The size of the patches was the same as the receptive field of the FCN.

The FCN consists of six convolutional blocks (Table S1). The first four convolutional blocks consist of a 3D convolutional layer followed by the operations: 3D max pooling, 3D batch-normalization, Leaky Relu and Dropout. The last two convolutional layers function as dense layers in terms of the classification task and these two layers play a key role in boosting model efficiency (*25*). The network was trained *de novo* with random initialization of the weights. We used the Adam optimizer with a 0.0001 learning rate and a mini-batch size of 10. During the training process, the model was saved when it achieved the lowest error on the ADNI validation dataset. After FCN training, a single volumetric MRI scan was forwarded to get a complete disease probability map (DPM) was an instantaneous process taking about a second on an NVIDIA GTX Titan GPU.

The FCN was trained by repeated application to cuboidal patches of voxels randomly-sampled from a full volume of sequential MRI slices. Since the convolutions decrease the size of the input across successive layers of the network, the size of each patch was selected such that the shape of the final output from each patch was equal to 2×1×1×1 (Table S1); i.e., the application of the FCN to each patch during training produced a list of two scalar values. These values can be converted to respective AD and NC probabilities by application of a softmax function, and the greater of the two probabilities was then used for classification of disease status. In this way, the model was trained to infer local patterns of cerebral structure that suggested an overall disease state.

After generating DPMs for all subjects, an MLP model framework was developed to perform binary classification to predict AD status by selecting AD probability values from the DPMs. This selection was based on observation of the overall performance of the FCN classifier as estimated using the MCC values on the ADNI training data. Specifically, we selected DPM voxels from 106 fixed locations that were indicated to have high MCC values (≥0.6). The features extracted from these locations served as input to the MLP model that performed binary classification of AD status (Model A in Step 3 in Fig. 1). Two additional MLP models were developed where one model used age, gender and MMSE score values as input to predict AD status (Model B in Step 3 in Fig. 1), and the other MLP took the 106 features along with age, gender and MMSE score as input to predict AD status (Model C in Step 3 in Fig. 1). All the MLP models comprised of a single hidden layer with width 64 and an output layer with size 2. The MLP models also included nonlinear operators such as ReLu and Dropout.

### Volumetric MRI segmentation and neuropathological validation

Cortical and sub-cortical structures from volumetric MRI scans of 11 individuals from the FHS cohort, with brain autopsies, were segmented using FreeSurfer (*28*). In-built functions such as ‘recon-all’, ‘mri_annotation2label’, ‘tkregister2’, ‘mri_label2vol’, ‘mri_convert’ and ‘mris_calc’ were used to obtain the segmented structures.

We validated the FCN model’s ability to identify regions of high AD risk by overlapping the predicted brain regions with postmortem findings. Eleven individuals from the FHS dataset had histopathological evaluations of autopsied brains, and 4 individuals out of the 11 had confirmed AD. A blinded assessment to all demographic and clinical information was conducted during the neuropathological evaluation. Detailed descriptions of the neuropathological evaluation have been previously reported (*29*). For this study, we examined the density of neurofibrillary tangles (NFTs), diffuse senile (DP), neuritic or compacted senile (NPL) plaques, from paraffin-embedded sections extracted within the cortical and sub-cortical regions. The sections were stained using Bielschowsky silver stain. Immunocytochemistry was performed for phosphorylated tau protein (Innogenetics, AT8, 1:2000) and amyloid-ß protein (Dako, 6F-3D, 1:500, pretreated in 90% formic acid for 2 minutes). The maximum density of NFTs per 200× field was assessed semi-quantitatively and scores ranging from 1-4 were assigned (1+: 1 NFT/field; 2+: 2-5 NFT/field; 3+: 6-9/field; and 4+: ≥10 NFT/field). Similarly, DP and NPL were examined in a 100× microscopic field and rated separately with scores ranging between 1-4 (1+: 1-9 plaques/field; 2+: 10-19/field, 3+: 20-32/field, and 4+: >32/field). The final determinations were made by averaging the count in 3 microscopic fields. The density of NFTs, NPLs and DPs in each brain region were qualitatively compared with the model’s AD probability in that region.

### Clinical validation

Nine US board-certified practicing neurologists and two non-US practicing neurologists (all referred to as neurologists) were asked to provide a diagnostic impression (AD versus NC) of 80 randomly selected cases from the ADNI dataset that were not used for model training. For each case, the neurologists were provided with full volumetric, T1-weighted MRI scan, subject’s age, gender and their MMSE score for evaluation. The same parameters were used for training the model (Model C in Fig. 1). To obtain estimates of how the deep learning model compared to an average neurologist, the characteristics of neurologist performance were averaged across the neurologists who individually evaluated each test case. More details on the neurologist approach to the ratings can be found in the Supplement.

### CNN model development

A three-dimensional (3D) convolutional neural network (CNN) was created to perform classification of AD and NC cases and compare its results with the FCN model. The CNN model was trained, validated and tested on the same split of data that was used for the FCN model. To facilitate direct comparison with the FCN model, one CNN model was developed using the MRI data alone, as well as an additional MLP which included the CNN model-derived features along with age, gender and MMSE score. Similar to the FCN-MLP model, we merged the CNN-based imaging features and non-imaging features at the dense layer.

The CNN model consisted of 10 convolutional layers followed by 3 dense layers (Fig. S6 & Table S6). Each convolution layer was followed by ReLu activations. Max-pooling layers between the convolution blocks were used to down-sample the feature maps. Nonlinear activation function, including ReLu, dropout, softmax and batch-normalization were applied on feature vectors of the dense layers. The CNN model was trained from scratch with the same optimizer and loss function as the FCN model. We used a learning rate of 0.0001 and mini-batch size of 6.

### Performance metrics

We first generated sensitivity-specificity (SS) and precision-recall (PR) curves based on model predictions on the ADNI test data as well as on the other independent datasets (NACC, AIBL and FHS). For each SS and PR curve, we also computed the area under curve (AUC) values. Additionally, we computed sensitivity, specificity, F1-score and Matthews correlation coefficient on each set of model predictions. The F1-score considers both precision and recall of a test and is defined as:

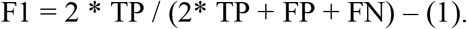

Here, TP denotes true positive values, and FP and FN denote false-positive and false-negative cases, respectively. MCC is a balanced measure of quality for dataset classes of different sizes of a binary classifier and defined as follows:

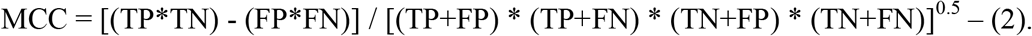

The, TN denotes true negative values. We also calculated inter-annotator agreement using Cohen’s kappa (κ), as the ratio of the number of times two annotators agreed on a diagnosis. The κ statistic measures interrater agreement for categorical items. A κ score of 1 indicates perfect agreement between the annotators. Average pairwise κ was computed that provided an overall measure of agreement between the neurologists.

### Statistical analysis

To assess the overall significant levels of differences between NC and AD groups, two-sample t-test and the Chi square test were used for continuous and categorical variables, respectively. The FCN model’s ability to identify regions of high AD risk was evaluated by overlapping the DPMs with postmortem histopathological findings. A subset of 11 individuals from the FHS study sample had undergone brain autopsy and were used for the analysis. In these participants, the locations and frequencies of Aβ and τ pathologies, semi-quantitatively reported by neuropathologists, were associated with high-AD risk regions. The Spearman’s rank correlation coefficient test was used to determine the strength and direction (negative or positive) of the relationship between these regional AD probabilities and pathology scores. Lastly, given the widespread recognition of diffuse cerebral atrophy in the normal aging process, we performed an age-based subgroup analysis. We used the ANOVA test to determine whether age distributions differed across TP, TN, FP and FN cases obtained from the multimodal MLP model (Model C). Separate analysis was conducted across each study cohort.

## Supplementary figures

**Figure S1:**
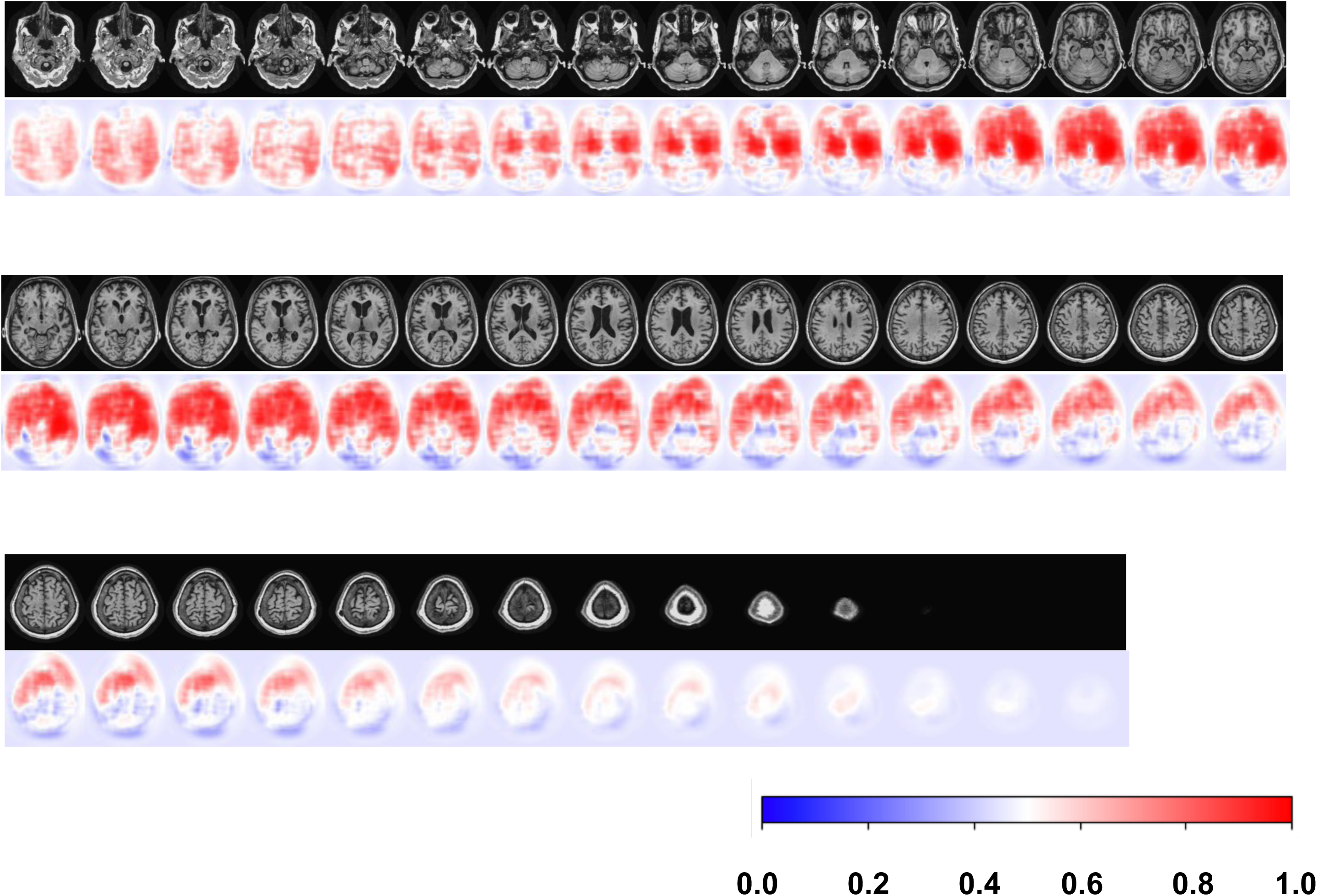
Axial stack of DPMs from a single subject. Local probabilities of disease were predicted for all slices of MRI stacks from each individual meeting inclusion criterion. All imaging planes were used to construct 3D DPMs. Here, we demonstrate the results of the model as applied to a full MRI sequence of an individual as viewed in the axial plane. Red color indicates locally-inferred probability of AD greater than 0.5, whereas blue indicates less than 0.5.

**Figure S2:**
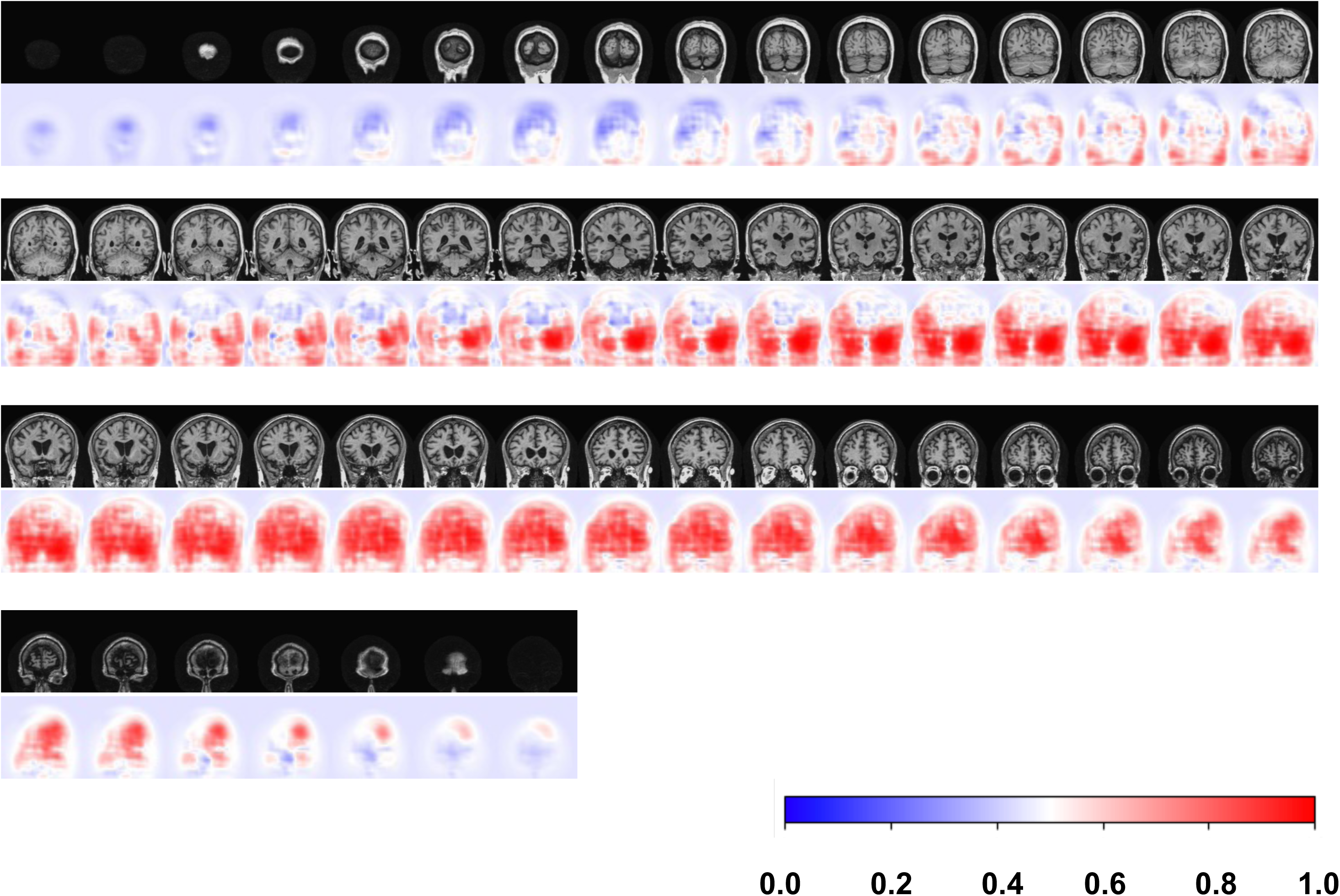
Coronal stack of DPMs from a single subject. Local probabilities of disease were predicted for all slices of MRI stacks from each individual meeting inclusion criterion. All imaging planes were used to construct 3D DPMs. Here, we demonstrate the results of the model as applied to a full MRI sequence of an individual as viewed in the coronal plane. Red color indicates locally-inferred probability of AD greater than 0.5, whereas blue indicates less than 0.5.

**Figure S3:**
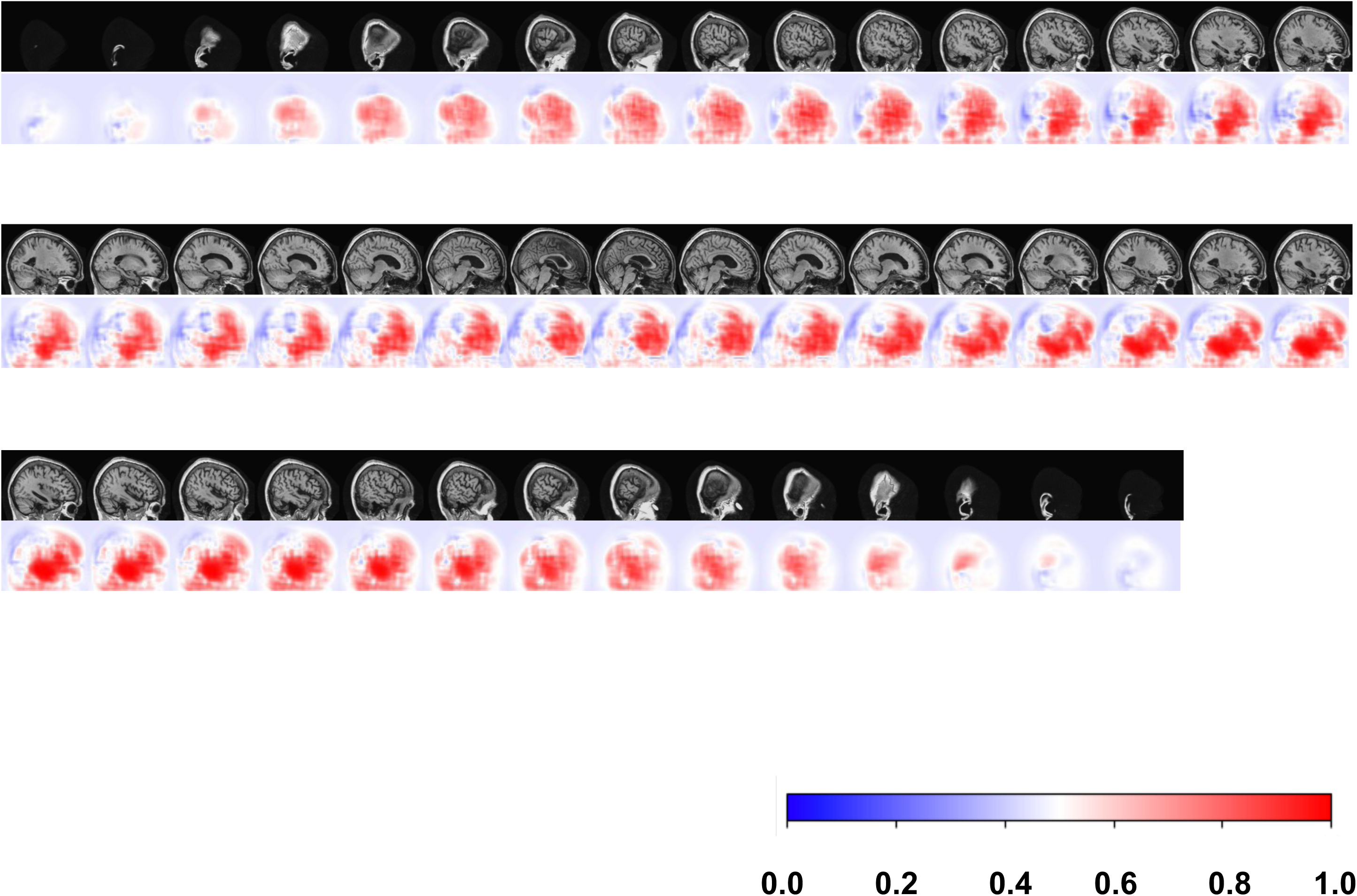
Sagittal stack of DPMs from a single subject. Local probabilities of disease were predicted for all slices of MRI stacks from each individual meeting inclusion criterion. All imaging planes were used to construct 3D DPMs. Here, we demonstrate the results of the model as applied to a full MRI sequence of an individual as viewed in the sagittal plane. Red color indicates locally-inferred probability of AD greater than 0.5, whereas blue indicates less than 0.5.

**Figure S4:**
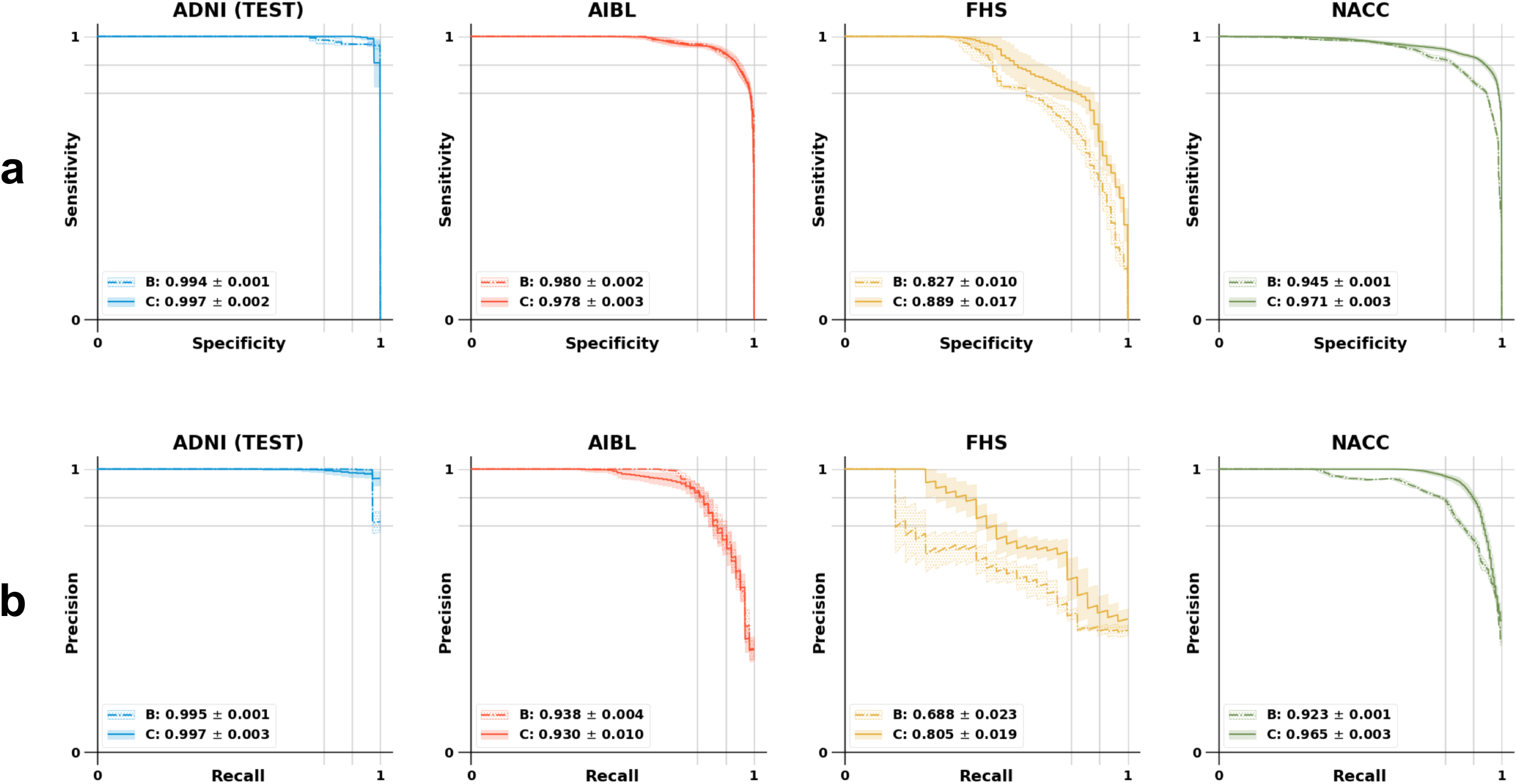
Performance of non-imaging and multimodal models utilizing ApoE status. ApoE status was added as an additional feature to the set of clinical variables utilized in the construction of the MLP and multimodal models described summarized in Figure 3a/3b. In this figure, Model B refers to the MLP model using age, gender, MMSE score and ApoE status as features whereas Model C refers to the MLP model that was developed using the FCN model features, age, gender, MMSE score and the ApoE status.

**Figure S5:**
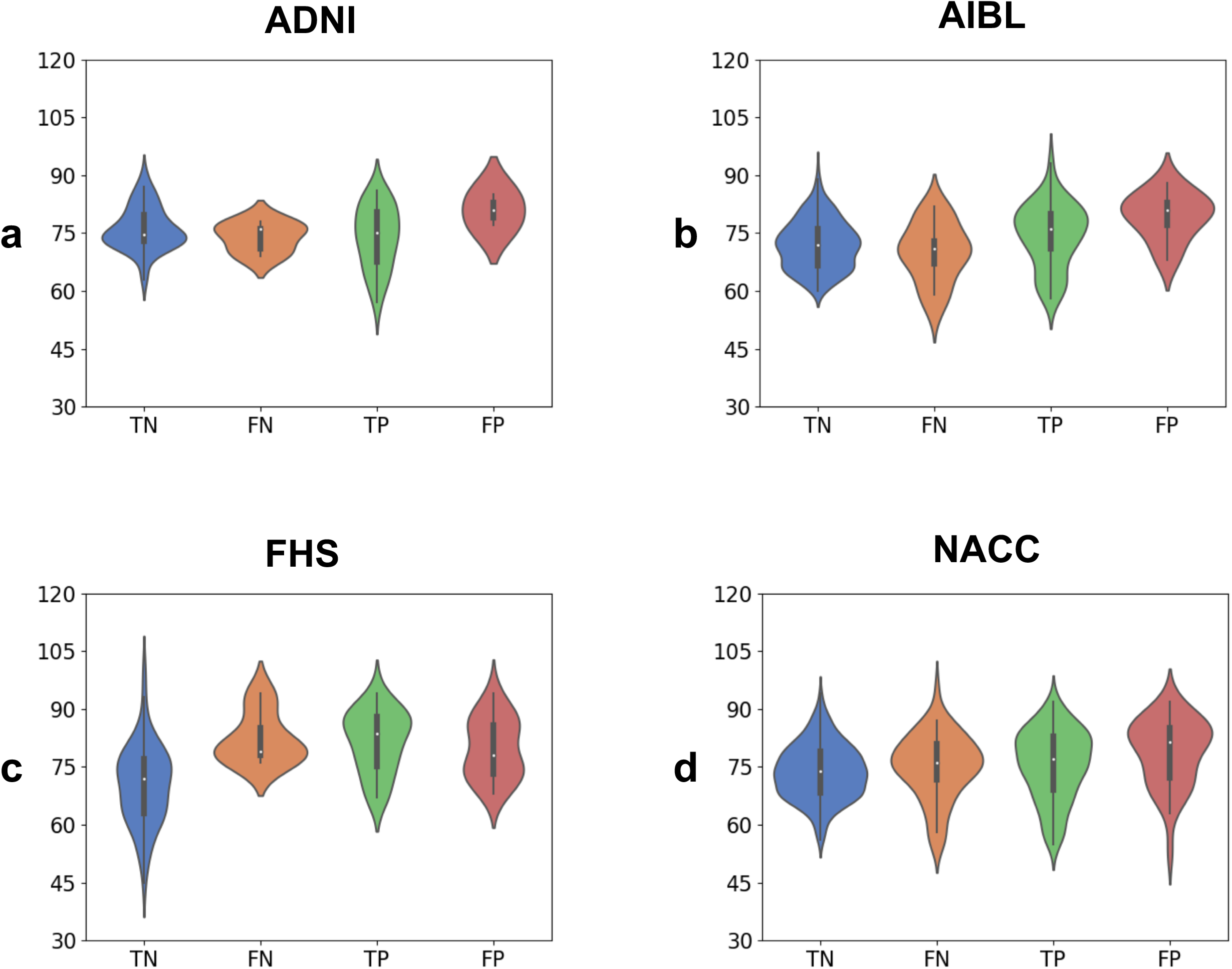
Violin plots of model performance by age. We conducted subgroup analyses on each dataset to determine potential effects of age in biasing the distribution of true negative, true positive, false negative, and false positive cases derived from the multimodal MLP model (Model C). This was particularly important given widespread recognition of diffuse cerebral atrophy in the normal aging process. Age was not found to have a large confounding effect on predictive performance.

**Figure S6:**
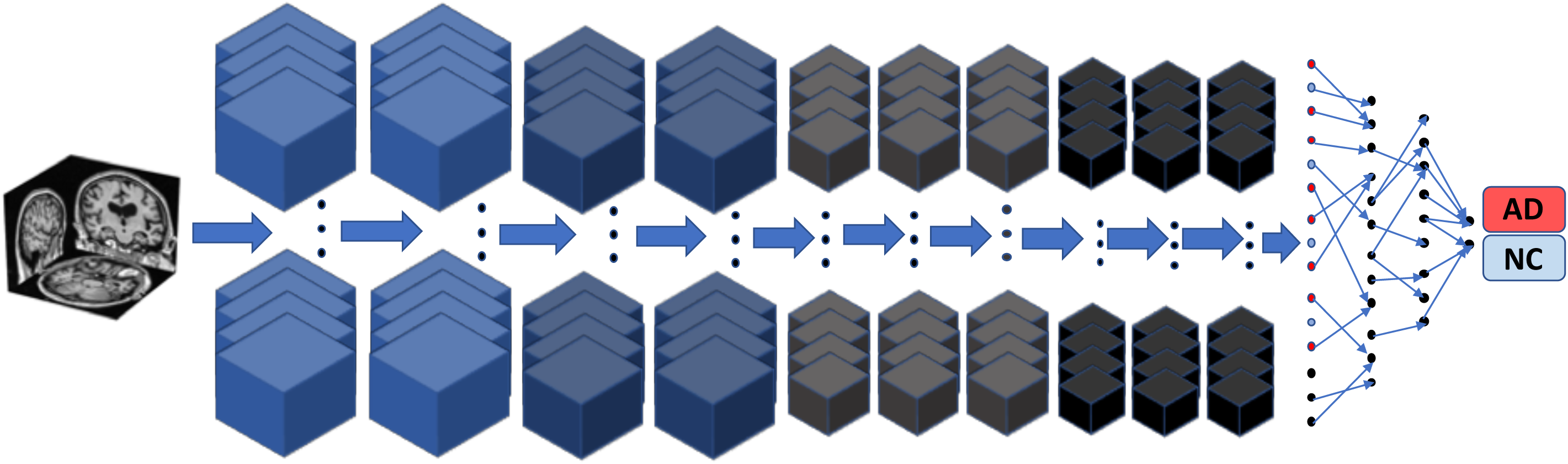
Schematic of the 3D convolutional neural network (CNN). The CNN model comprising 10 convolutional layers followed by 3 dense layers, was trained on the whole MRI volume to predict AD status.

**Figure S7:**
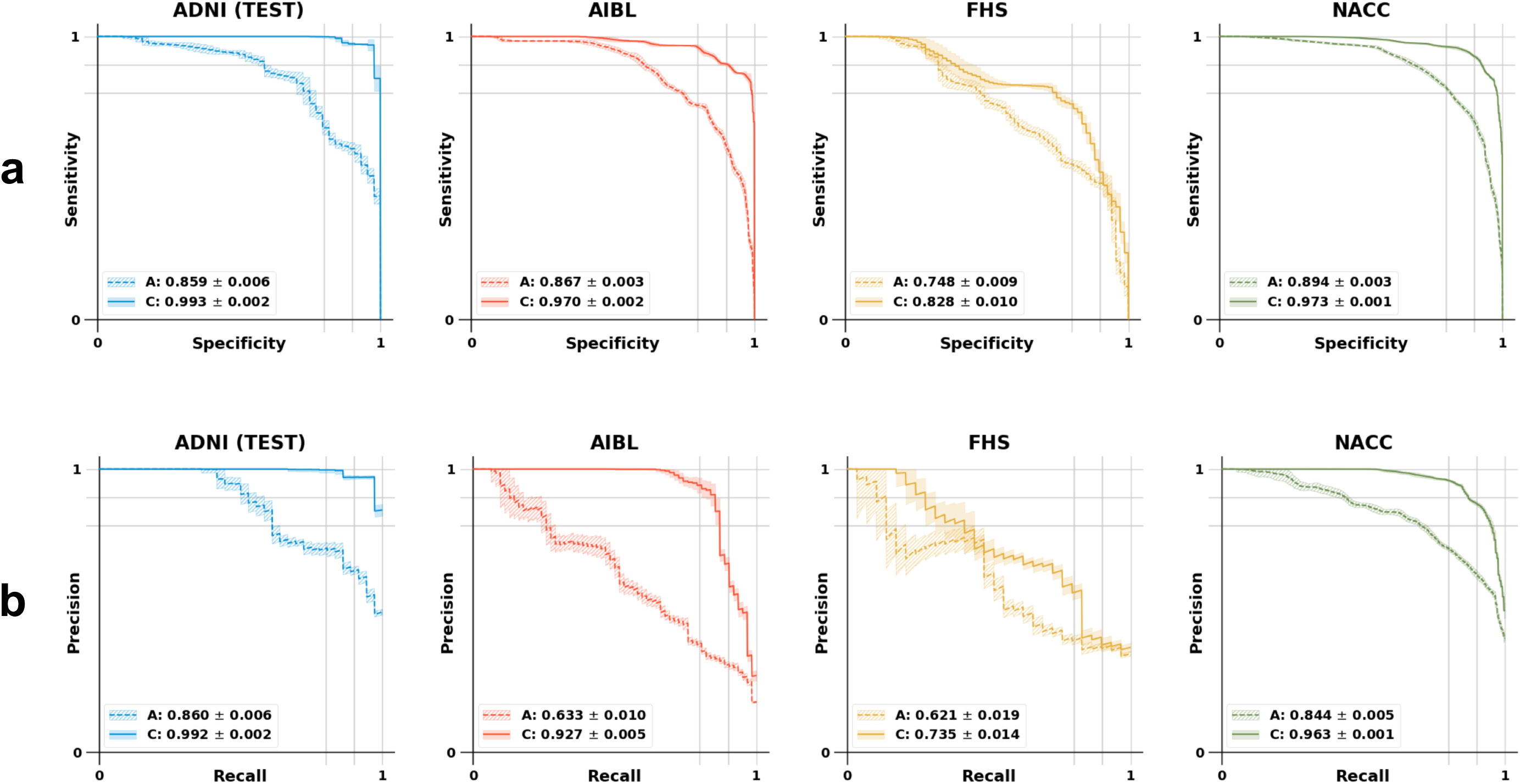
Performance of the 3D convolutional neural network (CNN). (a) SS curves comparing the CNN model developed using MRI as the sole input (Model A), and the other that included the CNN features and the non-imaging features including age, gender and MMSE score (Model C). (b) PR curves comparing the CNN model developed using MRI as the sole input (Model A), and the other that included the CNN features and the non-imaging features including age, gender and MMSE score (Model C).

## Supplementary tables

**Table S1:**
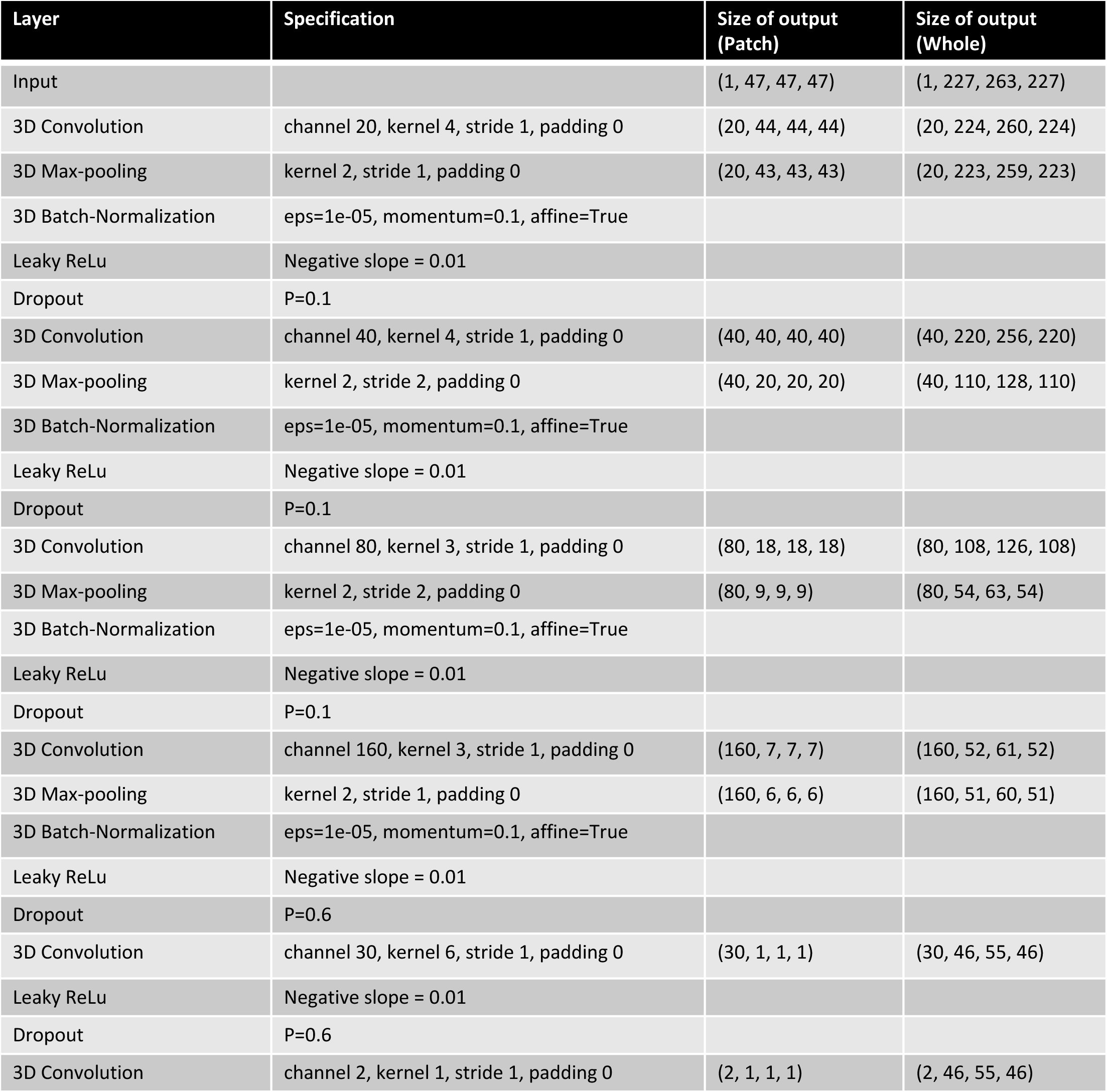
Summary of FCN architecture and hyperparameters for patch-wise training and full-volume application. The FCN model was trained on patches of size 47×47×47 in order to yield scalar (1×1×1) predictions of AD status from randomly-sampled sub-volumes. Each convolutional step within the network was followed by max-pooling and batch-normalization prior to passage to a leaky rectified linear unit (ReLu) activation. Channel depth, kernel size, padding, and stride hyperparameters are shown along with dropout probabilities at each step of the network. Application of the same model architecture to full-sized images yielded rank 3 tensors of size 46×55×46 that could be translated to DPMs via passage to a softmax function.

**Table S2:**
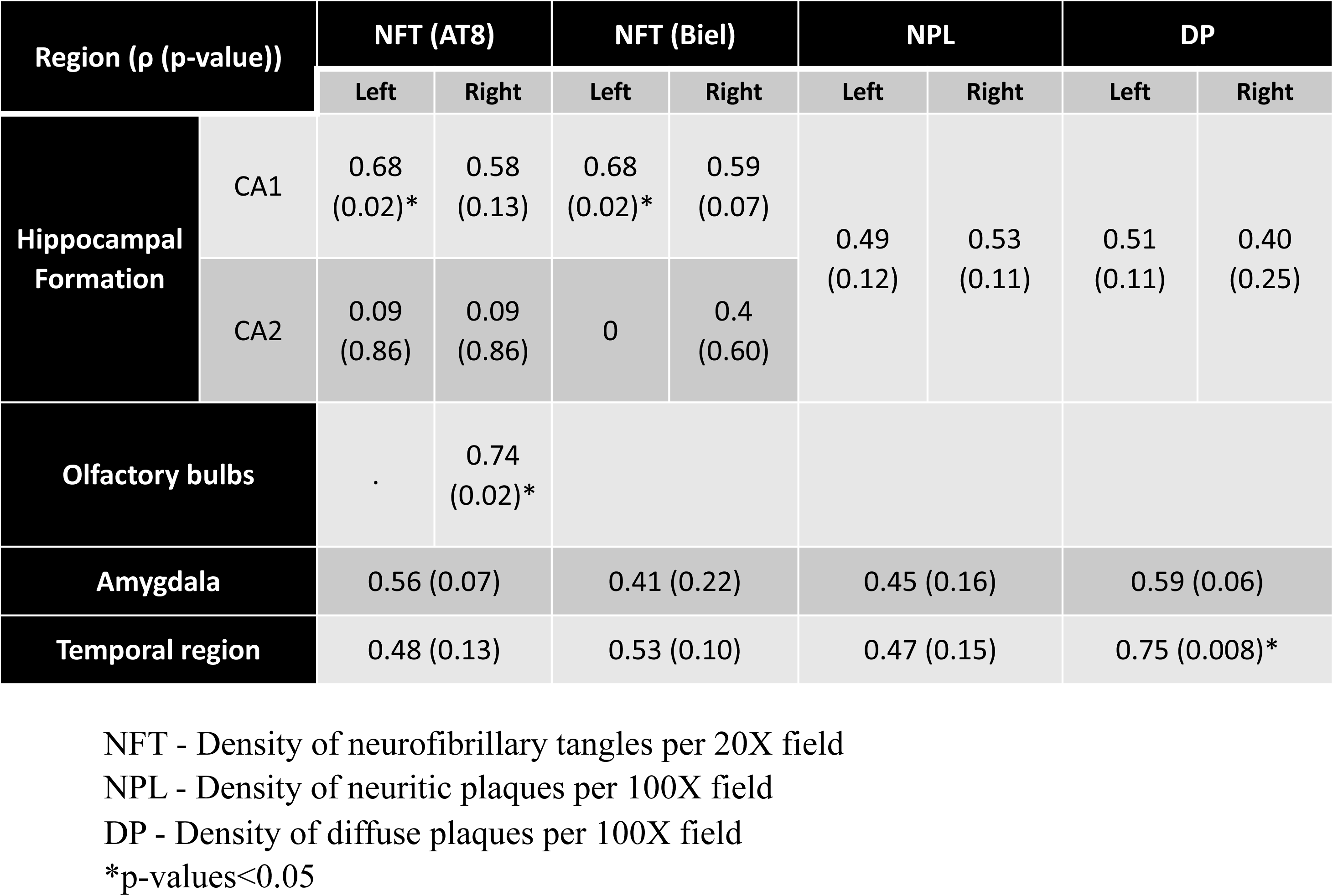
Correlation between neuropathologic findings and regional AD probability predictions. The MRI scans were segmented to represent various brain lobes using a segmentation platform (FreeSurfer), and population-averaged regional AD probabilities were computed for each segmented region. Spearman correlation coefficients were then calculated to quantify the relationship between the regional probabilities and semiquantitative pathology scores derived from pathologist-scored densities of neurofibrillary tangles (NFT), neuritic plaques (NPL) and diffuse plaques (DP). Using this non-parametric test, the strength and direction (negative or positive) of a relationship between the regional AD probabilities and pathology scores were assessed for significance at the 0.05 level. A positive correlation was observed across the numerous pathologies across these regions, however only the NFTs in the left hippocampal CA1 region, NFTs in the right olfactory bulb, and diffuse plaques in the temporal region reached statistical significance.

| Region (ρ (p-value)) |  | NFT (AT8) |  | NFT (Biel) |  | NPL |  | DP |  |
| --- | --- | --- | --- | --- | --- | --- | --- | --- | --- |
|  |  | Left | Right | Left | Right | Left | Right | Left | Right |
| Hippocampal Formation | CA1 | 0.68<br>(0.02)* | 0.58<br>(0.13) | 0.68<br>(0.02)* | 0.59<br>(0.07) | 0.49<br>(0.12) | 0.53<br>(0.11) | 0.51<br>(0.11) | 0.40<br>(0.25) |
|  | CA2 | 0.09<br>(0.86) | 0.09<br>(0.86) | 0 | 0.4<br>(0.60) |  |  |  |  |
| Olfactory bulbs |  | . | 0.74<br>(0.02)* |  |  |  |  |  |  |
| Amygdala |  | 0.56 (0.07) |  | 0.41 (0.22) |  | 0.45 (0.16) |  | 0.59 (0.06) |  |
| Temporal region |  | 0.48 (0.13) |  | 0.53 (0.10) |  | 0.47 (0.15) |  | 0.75 (0.008)* |  |

NFT - Density of neurofibrillary tangles per 20X field NPL - Density of neuritic plaques per 100X field DP - Density of diffuse plaques per 100X field \*p-values<0.05

**Table S3:** Summary of the models with ApoE included. Accuracy, sensitivity, specificity, F1-score and Matthew’s correlation coefficient (MCC) are displayed for the respective ApoE-inclusive models. Here, Model B refers to the MLP alone whereas Model C refers to the MLP model that was developed using the CNN model features, age, gender and MMSE score.

| Model B | Accuracy | Sensitivity | Specificity | F1-score | MCC |
| --- | --- | --- | --- | --- | --- |
| ADNI Test | 0.974±0.015 | 0.949±0.031 | 0.994±0.016 | 0.970±0.017 | 0.948±0.028 |
| AIBL | 0.940±0.016 | 0.881±0.036 | 0.951±0.024 | 0.828±0.032 | 0.796±0.036 |
| FHS | 0.752±0.030 | 0.706±0.056 | 0.771±0.054 | 0.627±0.030 | 0.454±0.048 |
| NACC | 0.862±0.012 | 0.884±0.025 | 0.849±0.031 | 0.822±0.009 | 0.716±0.016 |

b
| Model C | Accuracy | Sensitivity | Specificity | F1-score | MCC |
| --- | --- | --- | --- | --- | --- |
| ADNI Test | 0.966±0.013 | 0.942±0.027 | 0.986±0.011 | 0.962±0.015 | 0.933±0.025 |
| AIBL | 0.948±0.006 | 0.852±0.017 | 0.967±0.007 | 0.843±0.016 | 0.812±0.019 |
| FHS | 0.815±0.024 | 0.801±0.044 | 0.821±0.043 | 0.719±0.025 | 0.593±0.038 |
| NACC | 0.916±0.012 | 0.922±0.011 | 0.913±0.022 | 0.888±0.013 | 0.824±0.021 |

**Table S4:** Summary of FCN-based model performance. Three models were constructed for explicit performance comparison. Model A predicted AD status based upon imaging features derived from the patch-wise trained FCN. Model B consisted of a multilayer perceptron (MLP) that processed non-imaging clinical variables (age, gender, MMSE). Model C appended the clinical variables used by Model A to the MLP portion of Model B in order to form a multimodal imaging/non-imaging input. Accuracy, sensitivity, specificity, F1-score, and Matthew’s Correlation Coefficient (MCC) are demonstrated for each. Model C (multimodal) was found to outperform A and B in nearly all metrics in each of the four datasets. Of interest, however, we noted that the performance of Models A and B still displayed higher specificity and sensitivity than many of the human neurologists, all of whom used the full suite of available data sources to arrive at an impression (see Table S7 below).

| a | Model A | Accuracy | Sensitivity | Specificity | F1-score | MCC |
| --- | --- | --- | --- | --- | --- | --- |
|  | ADNI Test | 0.858±0.016 | 0.749±0.035 | 0.946±0.017 | 0.825±0.022 | 0.718±0.032 |
|  | AIBL | 0.893±0.004 | 0.641±0.026 | 0.941±0.006 | 0.660±0.013 | 0.597±0.015 |
|  | FHS | 0.828±0.016 | 0.815±0.072 | 0.833±0.019 | 0.740±0.035 | 0.620±0.048 |
|  | NACC | 0.852±0.005 | 0.794±0.020 | 0.885±0.015 | 0.798±0.006 | 0.681±0.009 |
| b | Model B | Accuracy | Sensitivity | Specificity | F1-score | MCC |
|  | ADNI Test | 0.954±0.014 | 0.927±0.034 | 0.976±0.031 | 0.947±0.016 | 0.908±0.027 |
|  | AIBL | 0.908±0.024 | 0.880±0.042 | 0.913±0.037 | 0.759±0.040 | 0.718±0.042 |
|  | FHS | 0.731±0.027 | 0.747±0.046 | 0.724±0.050 | 0.627±0.024 | 0.441±0.038 |
|  | NACC | 0.851±0.020 | 0.892±0.022 | 0.827±0.042 | 0.816±0.017 | 0.700±0.030 |
| c | Model C | Accuracy | Sensitivity | Specificity | F1-score | MCC |
|  | ADNI Test | 0.963±0.013 | 0.928±0.025 | 0.992±0.011 | 0.957±0.015 | 0.926±0.025 |
|  | AIBL | 0.946±0.005 | 0.851±0.014 | 0.965±0.006 | 0.838±0.014 | 0.806±0.016 |
|  | FHS | 0.824±0.023 | 0.812±0.034 | 0.829±0.036 | 0.736±0.026 | 0.613±0.040 |
|  | NACC | 0.915±0.009 | 0.913±0.011 | 0.916±0.018 | 0.888±0.010 | 0.821±0.016 |

**Table S5:** Summary of neurologist performance. Neurologists were recruited to perform validation of the deep learning model’s predictive performance. Each physician was presented with clinical information from 80 randomly-selected individuals from the ADNI cohort whose disease status was masked. For each case, MMSE score, age, and gender were given. Volumetric MRIs were made available for examination using an open source platform (http://www.slicer.org). Each neurologist provided diagnostic impressions using the given materials and accuracy, sensitivity, specificity, F1-score, and MCC were calculated relative to the clinical diagnosis. Of note, participating neurologists were requested to explain their reasoning when tasked to predict AD status from collections of MRIs, age, gender, and MMSE score for a subset of individuals with masked disease diagnosis. While there was notable variation in the order in which imaging and non-imaging data were considered, the two forms of information were widely considered complementary. Focal atrophy of key cerebral regions (notably the hippocampus and temporal lobes) was considered in light of generalized age-appropriate atrophy, and imaging was broadly utilized to rule out competing etiologies of dementia such as frontotemporal degeneration and vascular disease. MMSE, often considered in the context of age, was widely employed as comparison to salient imaging features as well. Collectively, these perspectives speak to the importance of an integrated approach to AD diagnosis in which distinct forms of information are reconciled prior to an ultimate classification of disease status.

|  | Accuracy | Sensitivity | Specificity | F1-score | MCC |
| --- | --- | --- | --- | --- | --- |
| Neurologist 1 | 0.850 | 0.833 | 0.864 | 0.833 | 0.697 |
| Neurologist 2 | 0.675 | 0.361 | 0.932 | 0.500 | 0.364 |
| Neurologist 3 | 0.888 | 0.861 | 0.909 | 0.873 | 0.772 |
| Neurologist 4 | 0.938 | 0.917 | 0.955 | 0.930 | 0.874 |
| Neurologist 5 | 0.888 | 0.917 | 0.864 | 0.880 | 0.777 |
| Neurologist 6 | 0.660 | 0.556 | 0.636 | 0.556 | 0.192 |
| Neurologist 7 | 0.825 | 0.611 | 1.000 | 0.759 | 0.681 |
| Neurologist 8 | 0.925 | 0.944 | 0.909 | 0.919 | 0.850 |
| Neurologist 9 | 0.825 | 0.806 | 0.841 | 0.806 | 0.646 |
| Neurologist 10 | 0.838 | 0.750 | 0.909 | 0.806 | 0.673 |
| Neurologist 11 | 0.738 | 0.444 | 0.977 | 0.604 | 0.513 |

**Table S6:** Summary of the convolutional neural network (CNN) architecture and hyperparameters. Each convolutional layer within the CNN model was followed by a rectified linear unit (ReLu) activation. Specific settings of each layers, i.e., channel depth, kernel size, padding, stride, dropout rate and momentum are shown.

| Layer | Specification | Size of output (Whole) |
| --- | --- | --- |
| Input |  | (1, 181, 217, 181) |
| 3D Convolution, ReLu | channel 8, kernel 3, stride 1, padding 0 | (8, 179, 215, 179) |
| 3D Convolution, ReLu | channel 8, kernel 3, stride 1, padding 0 | (8, 177, 213, 177) |
| 3D Max-pooling | kernel 2, stride 2, padding 0 | (8, 88, 106, 88) |
| 3D Convolution, ReLu | channel 16, kernel 3, stride 1, padding 0 | (16, 86, 104, 86) |
| 3D Convolution, ReLu | channel 16, kernel 3, stride 1, padding 0 | (16, 84, 102, 84) |
| 3D Max-pooling | kernel 2, stride 2, padding 0 | (16, 42, 51, 42) |
| 3D Convolution, ReLu | channel 32, kernel 3, stride 1, padding 0 |  |
| 3D Convolution, ReLu | channel 32, kernel 3, stride 1, padding 0 | (32, 40, 49, 40) |
| 3D Convolution, ReLu | channel 32, kernel 3, stride 1, padding 0 | (32, 38, 47, 38) |
| 3D Max-pooling | kernel 2, stride 2, padding 0 | (32, 19, 23, 19) |
| 3D Convolution, ReLu | channel 64, kernel 3, stride 1, padding 0 | (64, 17, 21, 17) |
| 3D Convolution, ReLu | channel 64, kernel 3, stride 1, padding 0 | (64, 15, 19, 15) |
| 3D Convolution, ReLu | channel 64, kernel 3, stride 1, padding 0 | (64, 13, 17, 13) |
| 3D Max-pooling | kernel 2, stride 2, padding 0 | (64, 6, 8, 6) |
| Flatten |  | (18432) |
| Dense, ReLu |  | (128) |
| Batch Normalization | eps=1e-05, momentum=0.1, affine=True |  |
| Dropout | P=0.7 |  |
| Dense, ReLu |  | (64) |
| Dense |  | (2) |
| Softmax |  |  |

**Table S7:** Summary of the 3D convolutional neural network (CNN) model performance. Accuracy, sensitivity, specificity, F1-score and Matthew’s correlation coefficient (MCC) are displayed for the 3D CNN model. Here, Model A refers to the CNN model alone whereas Model C refers to the MLP model that was developed using the CNN model features, age, gender and MMSE score.

| Model A | Accuracy | Sensitivity | Specificity | F1-score | MCC |
| --- | --- | --- | --- | --- | --- |
| ADNI Test | 0.738±0.042 | 0.704±0.152 | 0.766±0.174 | 0.703±0.045 | 0.497±0.056 |
| AIBL | 0.790±0.088 | 0.710±0.135 | 0.805±0.128 | 0.535±0.050 | 0.451±0.052 |
| FHS | 0.687±0.076 | 0.597±0.149 | 0.726±0.165 | 0.532±0.057 | 0.328±0.067 |
| NACC | 0.804±0.035 | 0.712±0.122 | 0.858±0.106 | 0.726±0.043 | 0.591±0.040 |

b
| Model C | Accuracy | Sensitivity | Specificity | F1-score | MCC |
| --- | --- | --- | --- | --- | --- |
| ADNI Test | 0.946±0.023 | 0.939±0.050 | 0.952±0.048 | 0.940±0.024 | 0.895±0.040 |
| AIBL | 0.907±0.050 | 0.892±0.038 | 0.910±0.066 | 0.770±0.082 | 0.732±0.089 |
| FHS | 0.777±0.030 | 0.721±0.081 | 0.801±0.069 | 0.661±0.026 | 0.507±0.035 |
| NACC | 0.907±0.024 | 0.898±0.046 | 0.913±0.059 | 0.878±0.021 | 0.809±0.035 |

## Code availability

Code for the algorithm development, evaluation, and statistical analysis will be made open source with no restrictions at the time of publication.

## Data availability

Data for the algorithm development will be made open source at the time of publication.

## Acknowledgments

This project was supported in part by the National Center for Advancing Translational Sciences, National Institutes of Health, through BU-CTSI Grant (1UL1TR001430), a Scientist Development Grant (17SDG33670323) from the American Heart Association, and a Hariri Research Award from the Hariri Institute for Computing and Computational Science & Engineering at Boston University, Framingham Heart Study’s National Heart, Lung and Blood Institute contract (N01-HC-25195; HHSN268201500001I) and NIH grants (R56-AG062109, AG008122, R01-AG016495, and R01-AG033040). Additional support was provided by Boston University’s Affinity Research Collaboratives program and Boston University Alzheimer’s Disease Center (P30-AG013846). V.B.K. also thanks Dr. Sang Chin at Department of Computer Science, Boston University for providing feedback on the manuscript.

## Author contributions

V.B.K. conceived, designed and directed the entire study. S.Q. designed and developed the fully convolutional network and performed image processing. C.X., S.Q., and G.H.C. designed and constructed the multilayer perceptron network. S.Q., P.S.J., C.X. and G.H.C. generated the figures. P.S.J. and S.Q. performed the analysis on the neuropathology data. C.K., S.Q., P.S.J. and M.I.M. performed data extraction. A.S.J. and C.K. performed MRI segmentation. S.Q., P.S.J. and C.X. performed sub-group analyses. B.D., S.Z., M.K., Y.Z., Y.J.A., A.S., S.K., M-H.S-H., S.H.A., J.Y., and E.A.S., are the practicing neurologists who reviewed the cases. V.B.K. and M.I.M. wrote the manuscript. R.A. provided the clinical relevance context. All the authors reviewed and edited the manuscript as well as provided scientific input.

## References

1. P. Scheltens et al., Alzheimer’s disease. Lancet 388, 505–517 (2016).

2. G. M. McKhann et al., The diagnosis of dementia due to Alzheimer’s disease: recommendations from the National Institute on Aging-Alzheimer’s Association workgroups on diagnostic guidelines for Alzheimer’s disease. Alzheimers Dement 7, 263–269 (2011).

3. R. C. Petersen et al., Alzheimer’s Disease Neuroimaging Initiative (ADNI): clinical characterization. Neurology 74, 201–209 (2010).

4. K. A. Ellis et al., Addressing population aging and Alzheimer’s disease through the Australian imaging biomarkers and lifestyle study: collaboration with the Alzheimer’s Disease Neuroimaging Initiative. Alzheimers Dement 6, 291–296 (2010).

5. J. M. Massaro et al., Managing and analysing data from a large-scale study on Framingham Offspring relating brain structure to cognitive function. Stat Med 23, 351–367 (2004).

6. D. L. Beekly et al., The National Alzheimer’s Coordinating Center (NACC) Database: an Alzheimer disease database. Alzheimer Dis Assoc Disord 18, 270–277 (2004).

7. G. B. Frisoni, N. C. Fox, C. R. Jack, Jr., P. Scheltens, P. M. Thompson, The clinical use of structural MRI in Alzheimer disease. Nat Rev Neurol 6, 67–77 (2010).

8. L. Harper, F. Barkhof, P. Scheltens, J. M. Schott, N. C. Fox, An algorithmic approach to structural imaging in dementia. J Neurol Neurosurg Psychiatry 85, 692–698 (2014).

9. C. R. Jack, Jr. et al., Tracking pathophysiological processes in Alzheimer’s disease: an updated hypothetical model of dynamic biomarkers. Lancet Neurol 12, 207–216 (2013).

10. N. I. Bohnen, D. S. Djang, K. Herholz, Y. Anzai, S. Minoshima, Effectiveness and safety of 18F-FDG PET in the evaluation of dementia: a review of the recent literature. J Nucl Med 53, 59–71 (2012).

11. A. Nordberg, PET imaging of amyloid in Alzheimer’s disease. Lancet Neurol 3, 519–527 (2004).

12. R. Ossenkoppele, et al., Associations between tau, Abeta, and cortical thickness with cognition in Alzheimer disease. Neurology 92, e601–e612 (2019).

13. N. Mattsson et al., Predicting diagnosis and cognition with (18)F-AV-1451 tau PET and structural MRI in Alzheimer’s disease. Alzheimers Dement, (2019).

14. T. G. Beach, S. E. Monsell, L. E. Phillips, W. Kukull, Accuracy of the clinical diagnosis of Alzheimer disease at National Institute on Aging Alzheimer Disease Centers, 2005-2010. J Neuropathol Exp Neurol 71, 266–273 (2012).

15. J. L. Whitwell et al., Neuroimaging correlates of pathologically defined subtypes of Alzheimer’s disease: a case-control study. Lancet Neurol 11, 868–877 (2012).

16. F. Barkhof et al., The significance of medial temporal lobe atrophy: a postmortem MRI study in the very old. Neurology 69, 1521–1527 (2007).

17. C. A. Raji, O. L. Lopez, L. H. Kuller, O. T. Carmichael, J. T. Becker, Age, Alzheimer disease, and brain structure. Neurology 73, 1899–1905 (2009).

18. L. A. van de Pol et al., Hippocampal atrophy in Alzheimer disease: age matters. Neurology 66, 236–238 (2006).

19. E. J. Topol, High-performance medicine: the convergence of human and artificial intelligence. Nat Med 25, 44–56 (2019).

20. G. Hinton, Deep Learning-A Technology With the Potential to Transform Health Care. JAMA 320, 1101–1102 (2018).

21. Y. LeCun, Y. Bengio, G. Hinton, Deep learning. Nature 521, 436–444 (2015).

22. S. Qiu et al., Fusion of deep learning models of MRI scans, Mini-Mental State Examination, and logical memory test enhances diagnosis of mild cognitive impairment. Alzheimers Dement (Amst) 10, 737–749 (2018).

23. D. Castelvecchi, Can we open the black box of AI? Nature 538, 20–23 (2016).

24. N. Coudray et al., Classification and mutation prediction from non-small cell lung cancer histopathology images using deep learning. Nat Med 24, 1559–1567 (2018).

25. J. Long, E. Shelhamer, T. Darrell, Fully Convolutional Networks for Semantic Segmentation. Proc Cvpr Ieee, 3431–3440 (2015).

26. L. van der Maaten, G. Hinton, Visualizing Data using t-SNE. J Mach Learn Res 9, 2579–2605 (2008).

27. D. Lu et al., Multimodal and Multiscale Deep Neural Networks for the Early Diagnosis of Alzheimer’s Disease using structural MR and FDG-PET images. Sci Rep 8, 5697 (2018).

28. B. Fischl, FreeSurfer. Neuroimage 62, 774–781 (2012).

29. R. Au et al., The Framingham Brain Donation Program: neuropathology along the cognitive continuum. Curr Alzheimer Res 9, 673–686 (2012).

